# Micron-scale, liquid-liquid phase separation in ternary lipid membranes containing DPPE

**DOI:** 10.1101/2025.05.08.652930

**Authors:** Gunnar J. Goetz, Sasha Naomi, Angelique M. Madrigal, Catherine L. A. Chang, Caitlin E. Cornell, Sarah L. Keller

**Author notes:** Department of Psychiatry, University of Nevada, Reno, NV, 89557. Department of Bioengineering, University of California, Berkeley, CA, 94720.

## Abstract

Micron-scale, liquid-liquid phase separation occurs in membranes of living cells, with physiological consequences. To discover which lipids might support phase separation in cell membranes and how lipids might partition between phases, miscibility phase diagrams have been mapped for model membranes. Typically, model membranes are composed of ternary mixtures of a lipid with a high melting temperature, a lipid with a low melting temperature, and cholesterol. Phospholipids in ternary mixtures are chosen primarily to favor stable membranes (phosphatidylcholines and sphingomyelins) or add charge (phosphatidylglycerols and phosphatidylserines). A major class of phospholipids missing from experimental ternary diagrams has been the phosphatidylethanolamines (PEs). PE-lipids constitute up to 20 mol% of common biological membranes, where they influence protein function and facilitate membrane fusion. These biological effects are often attributed to PE’s smaller headgroup, which leads to higher monolayer spontaneous curvatures and higher melting temperatures. Taken alone, the higher melting points of saturated PE-lipids imply that liquid-liquid phase separation should persist to higher temperatures in membranes containing PE-lipids. Here, we tested that hypothesis by substituting a saturated PE-lipid (DPPE) for its corresponding PC-lipid (DPPC) in two well-studied ternary membranes (DOPC/DPPC/cholesterol and DiphyPC/DPPC/cholesterol). We used fluorescence microscopy to map full ternary phase diagrams for giant vesicles over a range of temperatures. Surprisingly, we found no micron-scale, liquid-liquid phase separation in vesicles of the first mixture (DOPC/DPPE/cholesterol), and only a small region of liquid-liquid phase separation in the second mixture (DiphyPC/DPPE/cholesterol). Instead, coexisting solid and liquid phases were widespread, with the solid phase enriched in DPPE. An unusual feature of these ternary membranes is that solid and liquid-ordered phases can be distinguished by fluorescence microscopy, so tie-line directions can be estimated throughout the phase diagram, and transition temperatures to the 3-phase region (containing a liquid-disordered phase, a liquid-ordered phase, and a solid phase) can be accurately measured.

**SIGNIFICANCE STATEMENT:** Under physiological conditions, yeast vacuole membranes phase separate into liquid phases. The resulting domains are microns in size, and they are important for the cell’s function. Yeast membranes contain a significant fraction of lipids with phosphatidylethanolamine headgroups. It was unknown whether these lipids enhanced liquid-liquid phase separation or hindered it. Here, we produced model membranes containing a saturated phosphatidylethanolamine lipid, we mapped miscibility phase diagrams over broad temperature ranges, and we compared our results to existing diagrams for membranes of other lipid types. We were surprised to find that the new lipid suppressed liquid-liquid phase separation compared to a lipid with a larger, phosphatidylcholine headgroup. Previous simulations incorporated phosphatidylethanolamine lipids, and our results provide an experimental basis for future simulations.

## INTRODUCTION

Liquid-liquid phase separation of membranes is key to the function of some cells, which adjust their membrane compositions with respect to the phase transition (1, 2). Independent of whether membranes are derived from a cell with hundreds of lipids (2–5), or from minimal model systems with only two lipids (6), all membranes that demix into micron-scale liquid phases contain at least three moieties: lipid chains with high order, lipid chains with low order, and sterols (7–9).

**Figure 1.**
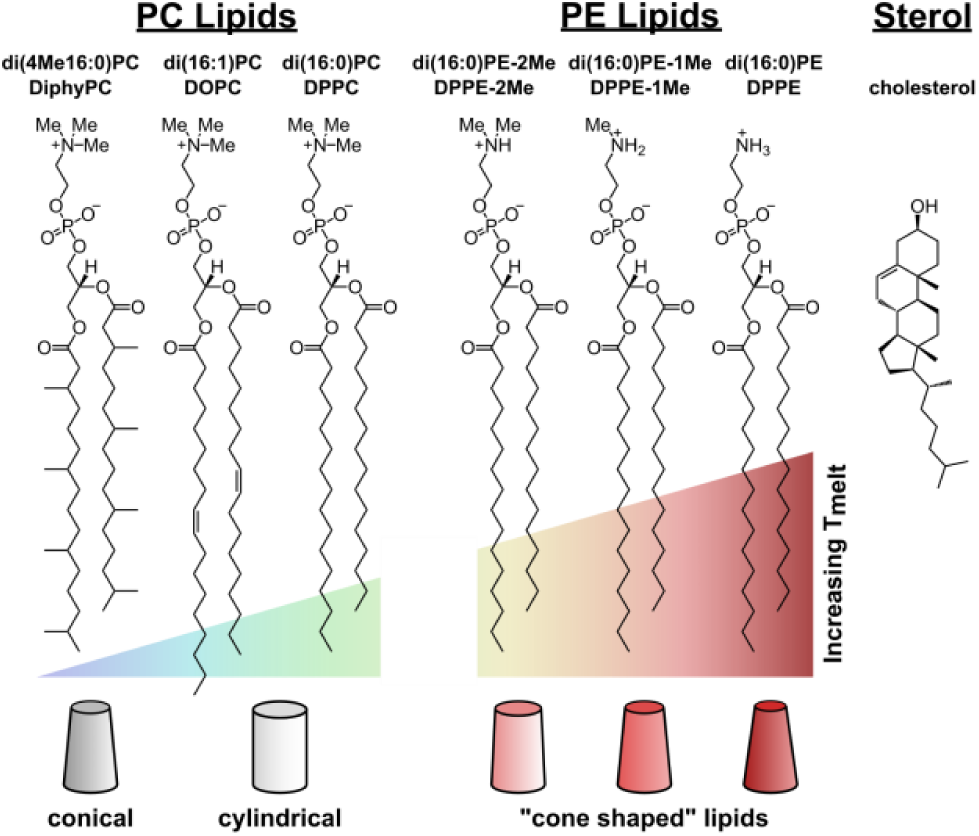
Structures of lipids. Phospholipids are ordered from left to right in terms of increasing melting temperature (*T*_melt_). Vesicles were made from ternary mixtures of lipids with a high *T*_melt_, lipids with a low *T*_melt_, and a sterol. Lipids have noncylindrical shapes when they have highly branched or unsaturated chains (as for DiPhyPC at the left) and/or smaller headgroups (as for PE lipids on the right). Headgroups of PC- and PE-lipids differ only by the number of methyl groups on the amine.

To discover which types of lipids support liquid-liquid phase separation and how the lipids partition between the phases, researchers have mapped phase diagrams, typically of ternary mixtures of a lipid with saturated chains, a lipid with either unsaturated or methylated chains, and cholesterol (Fig. 1). When researchers are faced with the question of exactly which lipids to incorporate into ternary membranes, they often choose PC-lipids, which are widely commercially available and produce stable membranes over broad experimental temperatures (10). Although researchers have made forays into producing ternary membranes containing other types of lipids (e.g., sphingomyelins (11, 12), ceramides (13, 14), and charged PS- or PG-lipids (15, 16)), mapping phase diagrams can be expensive, in both materials and time. As a result, there have been major gaps in the literature, for example in the lack of ternary membranes containing phosphatidylethanolamine (PE) lipids.

PE-lipids are distinguished by smaller headgroups than PC-lipids and by the potential to hydrogen bond with neighboring phosphates, both of which contribute to the cone shape of PE-lipids (17, 18). In turn, PE-lipids have higher chain order than corresponding PC-lipids, and they impart higher chain order in neighboring lipids (19). Higher chain order parameters hold whether they are assessed at equivalent absolute temperatures or at equivalent reduced temperatures (20). Higher chain order results in lower molecular areas (21) and higher *T*_melt_ values. For example, for di(16:0)PE, also known as DPPE, *T*_melt_ = 62.3 ± 5.0°C (22), whereas for di(16:0)PC, also known as DPPC, *T*_melt_ = 41.3 ± 1.8°C (10).

There are several ways experimentalists might use phase diagrams of membranes with PE-lipids. Knowledge of a membrane’s phase diagram would allow researchers to incorporate channels and proteins into membranes that undergo liquid-liquid phase separation such that local enrichment or depletion of PE-lipids in domains would enhance or diminish the effect of PE-lipids. This would allow researchers to leverage the tendency of some molecules to function and fold differently in membranes containing PE-lipids (e.g., (23–26), with other results reviewed in (27)). Beyond protein function, the spontaneous curvature that cone-shaped lipids impart to each leaflet of a membrane (or to the whole of an asymmetric membrane) has broad biological implications. Curvature can influence membrane fusion (28) as well as the size and shape of phase-separated domains ((29), and reviewed in (30)). Likewise, high fractions of PE-lipids are associated with transitions from flat, lamellar sheets to curved, inverted hexagonal (H_II_) and cubic phases (22, 31). The pressure dependence of the lamellar-H_II_ transition (32) explains why deep-sea creatures such as ctenophores undergo “tissue melting” at atmospheric pressure (33). Knowledge of which PE-containing membranes form stable vesicles under common laboratory conditions informs the types of experiments that researchers can design.

Theorists might use experimental phase diagrams of membranes with PE-lipids as ground truth for the models they create. Just as original phase diagrams for membranes with PC-lipids (34, 35) inspired a phenomenological model based on chain order parameters (36), phase diagrams for membranes with PE-lipids could aid in the development of new models that incorporate terms to account for PE headgroups. PE-lipids are common in biological membranes. They constitute ∼10% of red blood cell membranes (3), ∼8% of giant plasma membrane vesicles from zebrafish ZF4 cells (2), ∼15% of total yeast membranes (37), and 10-20% of yeast vacuole membranes (4, 5). As scientists incorporate more diverse lipids into their simulations (reviewed in (38)), phase diagrams that incorporate more headgroup types become more valuable, particularly in the context of curvature stress of asymmetric membranes (39).

Here, we produce giant unilamellar vesicles (GUVs) composed of ternary mixtures of cholesterol, a PE-lipid with a high melting temperature (DPPE, or di(16:0)PE), and a PC-lipid with a low melting temperature (either DOPC/di(18:1)PC or DiphyPC/di(16:0-4Me)PC). We chose DPPE as our PE-lipid because it has been reported to form a lamellar phase at common experimental temperatures (22). Even so, membranes with DPPE likely experience some curvature frustration because DPPE transitions into an inverted hexagonal phase at 120°C (22), and a cubic phase may lie nearby (32, 40). In our experiments, we use fluorescence microscopy to identify coexisting phases in vesicle membranes, and we map their miscibility phase diagrams. We compare our data with existing phase diagrams for membranes without PE-lipids, and we adjust the methylation of lipid headgroups to understand intermediate chemical structures between PE-lipids and PC-lipids.

## MATERIALS AND METHODS

### Lipids

1,2-dipalmitoyl-sn-glycero-3-phosphoethanolamine (DPPE or di(16:0)PE, *T*_melt_ = 62.3 ± 5.0°C, *T*_hex_ = 120.6 ± 3.6°C (22)), 1,2-dipalmitoyl-sn-glycero-3-phosphoethanolamine-N-methyl (DPPE-1Me or di(16:0)PE-1Me, *T*_melt_ = 57.6 ± 1.5°C (22)), 1,2-dipalmitoyl-sn-glycero-3-phosphoethanolamine-N,N-dimethyl (DPPE-2Me or di(16:0)PE-2Me, *T*_melt_ = 43.2 ± 2.0°C (22)), 1,2-dipalmitoyl-sn-glycero-3-phosphocholine (DPPC or di(16:0)PC, *T*_melt_ = 41.3 ± 1.8°C (10)), 1,2-dioleoyl-sn-glycero-3-phosphocholine (DOPC or di(16:1)PC, *T*_melt_ = -18.3 ± 3.6°C (10)), 1,2-diphytanoyl-sn-glycero-3-phosphocholine (DiPhyPC or di(16:0-4Me)PC, *T*_melt_ = <-120°C (41)), and a fluorescent dye, rhodamine-DPPE, were obtained from Avanti Polar Lipids (Alabaster, AL). Cholesterol was obtained from Sigma Aldrich (St. Louis, MO). All lipids were used without further purification. Cholesterol was mixed from powder in chloroform to a concentration of 10 mg/ml. DPPE, DPPE-1Me, and DPPE-2Me were mixed from powder in 9:1 chloroform/methanol to a concentration of 10 mg/ml. All other unlabeled lipids were supplied from Avanti in chloroform, and their nominal concentrations of 10 mg/ml were not verified. Lipid stocks were sealed with Teflon tape and parafilm and stored at -20°C.

### Electroformation

Vesicles were made by electroformation (42). Two halves of a glass slide coated with indium-tin-oxide (ITO) (Delta Technologies, Loveland, CO) were heated on a heat block at 60°C, which is just below the boiling point of chloroform. A comparison of data from ITO-coated slides on a heat block at 75°C appears in Fig. S1. Stocks of phospholipids and cholesterol (with 0.8 mol% rhodamine-DPPE) in chloroform or chloroform/methanol were mixed to achieve a total of 0.25 mg total lipid and heated to 60°C for 15 min on the heat block. The lipid solutions were spread evenly across the slides with the side of a glass Pasteur pipette. As the solvent evaporated from the lipid layer, thin-film interference was visible by eye. The slide-halves were exposed to vacuum for ≥30 min to allow any remaining chloroform to evaporate. The two halves were assembled face-to-face, separated by Teflon spacers of 0.3 mm thickness held in place with vacuum grease. The chamber was filled with 18 MΩ–cm water, sealed with vacuum grease, and attached to metal electrodes using binder clips. An AC voltage of 1.0 V at 10 Hz was applied across the electrodes for 1-2 hours at 75°C while vesicles formed (60°C for DiphyPC/DPPC/cholesterol). This temperature was chosen to be well above the highest *T*_melt_ of any lipid used, to adequately incorporate all lipids into vesicles and to minimize error in transition temperatures (43). For vesicles containing DPPE, 60°C is not a sufficiently high electroformation temperature (Fig. S2). After electroformation, vesicles were transferred into an Eppendorf tube and typically imaged immediately (or within two hours). From limited data on how lipid ratios in vesicles differ from ratios in stock solutions, we expect at least three of the lipids in our study to faithfully incorporate into electroformed vesicles based on data for DOPC, cholesterol, and a saturated PE-lipid (DLPE; di(12:0)PE) (44).

### Vesicle Imaging

Vesicle solutions were diluted to achieve roughly 100-200 vesicles per field of view (usually a 10:1 dilution) and placed between #1.5 glass coverslips. Coverslip edges were coated with vacuum grease, a second coverslip was placed on top to seal the vesicle solution inside, and the coverslip assembly was placed on a layer of Omegatherm thermal paste (Omega Engineering, Norwalk, CT) coating the top plate of a Peltier temperature controller on a home-built microscope stage. Vesicles were imaged on a Nikon (Melville, NY) Eclipse ME600L upright epifluorescence microscope using a Hamamatsu (Bridgewater, NJ) C13440 camera. Contrast between phases in the membrane arises from preferential partitioning of the lipid dye into the phase with less lipid order. For every sample containing membranes with coexisting phases, a representative image was selected for a field of view of vesicles at ∼30°C (Fig. S4-6 and S9-11). Fiji (45) was used to crop the image to 300 x 300 pixels, to adjust brightness and contrast so that vesicle features were visible (gamma = 1), and to add scalebars.

### Transition Temperatures and Plotting

We use *T*_mix_ to denote three possible transitions that result in an increase in the number of phases in a membrane as the temperature decreases: L→ L_o_/L_d_, L→ S/L, and S/L→ S/L_o_/L_d_. The corresponding phase transition temperatures were first estimated, then measured. To make estimates, an initial coverslip assembly was heated at ∼1°C/min, and the approximate range of temperatures over which phase-separated domains disappeared was recorded. Temperature was then set 5-10°C above the estimated transition, and the initial coverslip assembly was discarded and replaced with a new one. Fields of vesicles (usually >100 vesicles total) were imaged as temperature was decreased stepwise over a total span of ≥12°C. At each step, temperature was allowed to equilibrate for ∼1-2 min before images were collected. For each field of view, the number of vesicles with phase-separated membranes and the number with uniform membranes were tabulated by hand. Circular domains that immediately coalesced upon collision were categorized as liquid phase. Solid phases were categorized as having domains with irregular shapes and/or domains that did not coalesce upon collision. In vesicles with very small area fractions of ordered phase, domains are so small that L_o_ and solid phases are difficult to distinguish. To determine the transition temperature, *T*_mix_, a non-linear least squares algorithm from the open-source Python library, SciPy (46), was used to fit a sigmoidal curve of the form

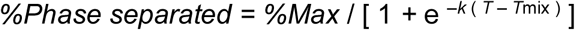

where %*Max* is the maximum percent of vesicles that phase separate, *k* is the steepness of the curve, *T_mix_*is the phase transition temperature located at the inflection point, and *T* is the temperature input variable. For all samples, *%Max* ≥ 90%. Plots of sigmoidal curves and fits were generated using the open-source Python library, Matplotlib (47), and are shown in Fig. S13-18. Plots of ternary phase diagrams were generated using the open-source Python library, mpltern (48). Interpolation between data points was achieved with the griddata function in SciPy (46), which uses a Clough-Tocher scheme. All phase transition temperatures and errors are recorded in Tables S1-S8.

### Measurement Errors

Error was quantified in four ways: **1)** For each sigmoidal curve, a 95% confidence interval (shown in Fig. S13-S18) was calculated, which provides errors of the fit parameters. For *T*_mix_, these fit errors are very small (<0.5°C) and are recorded in Tables S1-S8. **2)** For each sigmoidal curve, ½(|*T*_90%_ *– T*_10%_|) was calculated; this is half the range between the temperatures at which 90% of vesicles and 10% of vesicles are phase separated (Fig. S13-S18). These errors are recorded in Tables S1-S8. **3)** Three researchers tabulated numbers of phase-separated vesicles. To estimate the error that each individual introduced, all three researchers tabulated the same four data sets in Fig. S19. The resulting values of *T*_mix_ were typically within ±0.5°C. **4)** Sample-to-sample variation was measured by standard deviations across three duplicate sample sets and nineteen triplicate sample sets, shown in bolded data in Tables S1-8. The average standard deviation across all duplicate and triplicate sample sets was 0.7 ± 0.6°C and ranged from 0.1°C to 2.8°C.

## RESULTS AND DISCUSSION

### Vesicles are stable over wide ranges of DPPE mole fractions

Ternary lipid mixtures containing DPPE produce stable, giant vesicles over broad spans of temperature and mole fraction (Fig. 2). The solubility of cholesterol in PE-membranes is lower than in equivalent PC-membranes – we maintained cholesterol fractions at ≤50 mol% to avoid exceeding cholesterol’s solubility limit of 51 ± 3 mol% in DPPE membranes (49, 50). Although DPPE’s saturated chains make it less conical than unsaturated PE-lipids, challenges persist in forming ternary vesicles with DPPE. In ternary mixtures, when the combined mole fraction of DPPE and cholesterol exceeds 80-90 mol%, yields of giant vesicles are too low for data collection. In binary mixtures, yields of vesicles electroformed at 75°C are low at ≥60 mol% DPPE (Fig. 2), consistent with vesicles made by gentle hydration in water at 70°C (51). No significant difference in membrane composition is expected for vesicles made by gentle hydration versus electroformation (44). Two other groups reported the formation of binary vesicles at higher DPPE concentrations by gentle hydration but did not report the temperature of hydration (52, 53). Specifically, Blume and Griffin evaporated solvent at 60°C, hydrated, and held vesicles at 60°C for 1 hour before collecting data (52), whereas Wu and McConnell hydrated lipid films in 100 µM sodium phosphate buffer at an unspecified temperature above the lipids’ upper transition (53). Vesicles produced at lower temperatures (e.g., 60°C rather than 75°C) appear to incorporate less DPPE than intended (Fig. S2).

**Figure 2.**
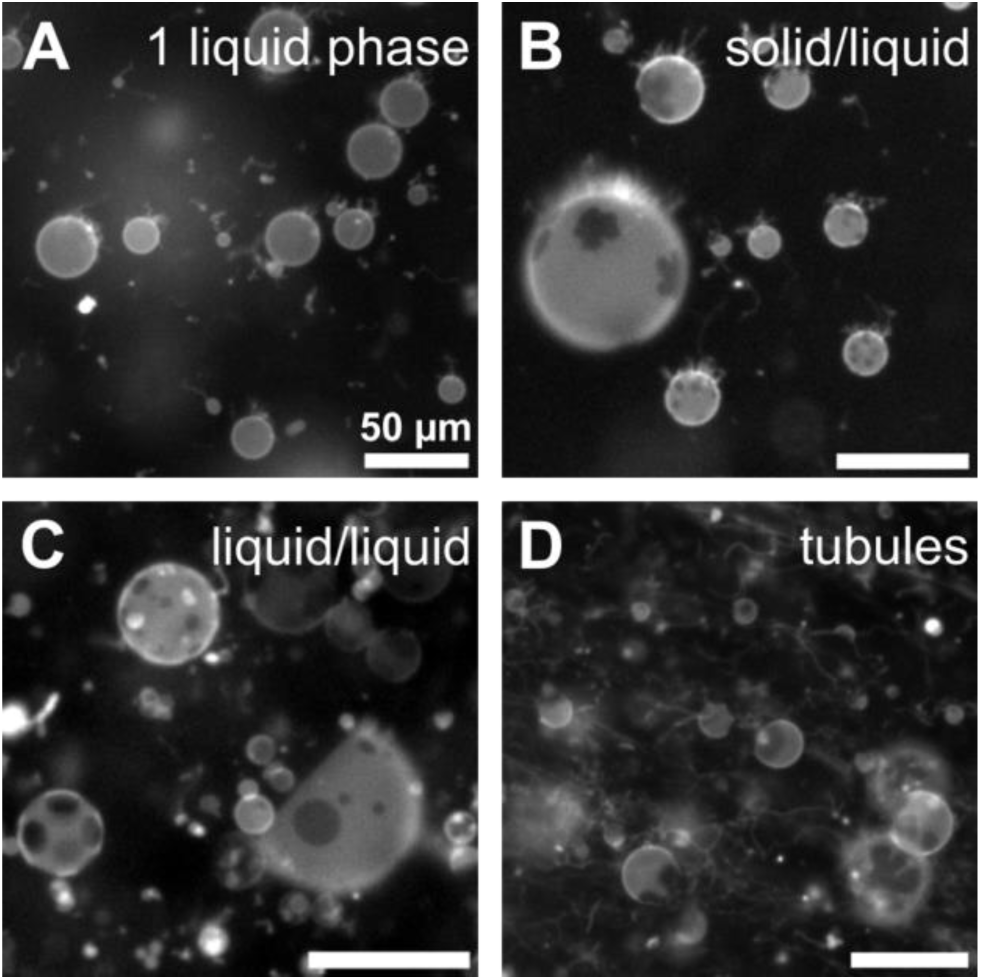
Lipid mixtures containing DPPE can form stable vesicles that phase separate, often with tubes. A) At high temperature (50°C), vesicle membranes of 20:50:30 DOPC/DPPE/cholesterol are in one liquid phase. B) At lower temperature (46°C), vesicles in a different field of view of the same sample phase separate into coexisting solid (dark) and liquid (bright) phases. Solid phases are characterized by noncircular domains that do not coalesce over time. C) Coexisting L_o_ and L_d_ liquid phases appear for some ratios of DiphyPC/DPPE/cholesterol, shown here for 30:40:30 at 45°C. Liquid phases are characterized by circular domains that merge in taut vesicles. D) A “worst case scenario” of long tubules emanating from vesicles containing DPPE, shown here for a ratio of 10:60:30 of DOPC/DPPE/cholesterol at 52°C. Contrast between the two phases arises from 0.8 mol % fluorescently labeled rhodamine-DPPE, which preferentially partitions to the L_d_ phase. More representative fields of view are shown in Figs. S4-6 and Figs. S8-11.

### Replacing DPPC with DPPE eradicates (or severely limits) liquid-liquid coexistence in ternary membranes

Ternary membranes containing DPPE differ starkly from membranes containing the corresponding PC-lipid. In Fig. 3, liquid-liquid coexistence is obliterated across the entire phase diagram of DOPC/DPPE/cholesterol, whereas it persists across broad ranges of temperature and lipid composition in vesicles of corresponding PC-lipids (Fig. 3C) (34, 54). In place of liquid-liquid coexistence, solid-liquid coexistence is observed (Fig. 3A-B). Solid domains are often identified by non-circular shapes, similar to the hexagonal and flower-shaped domains in binary DOPC/DPPC vesicles (55, 56). The area fraction of dark, noncircular, solid domains in vesicles monotonically increases as the mole fraction of DPPE increases (toward the bottom right vertex in the small images in Fig. 3A and in the large fields of view in Fig. S4-S6). This observation is in line with the expectation that solid phases are enriched in lipids with high melting temperatures.

**Figure 3.**
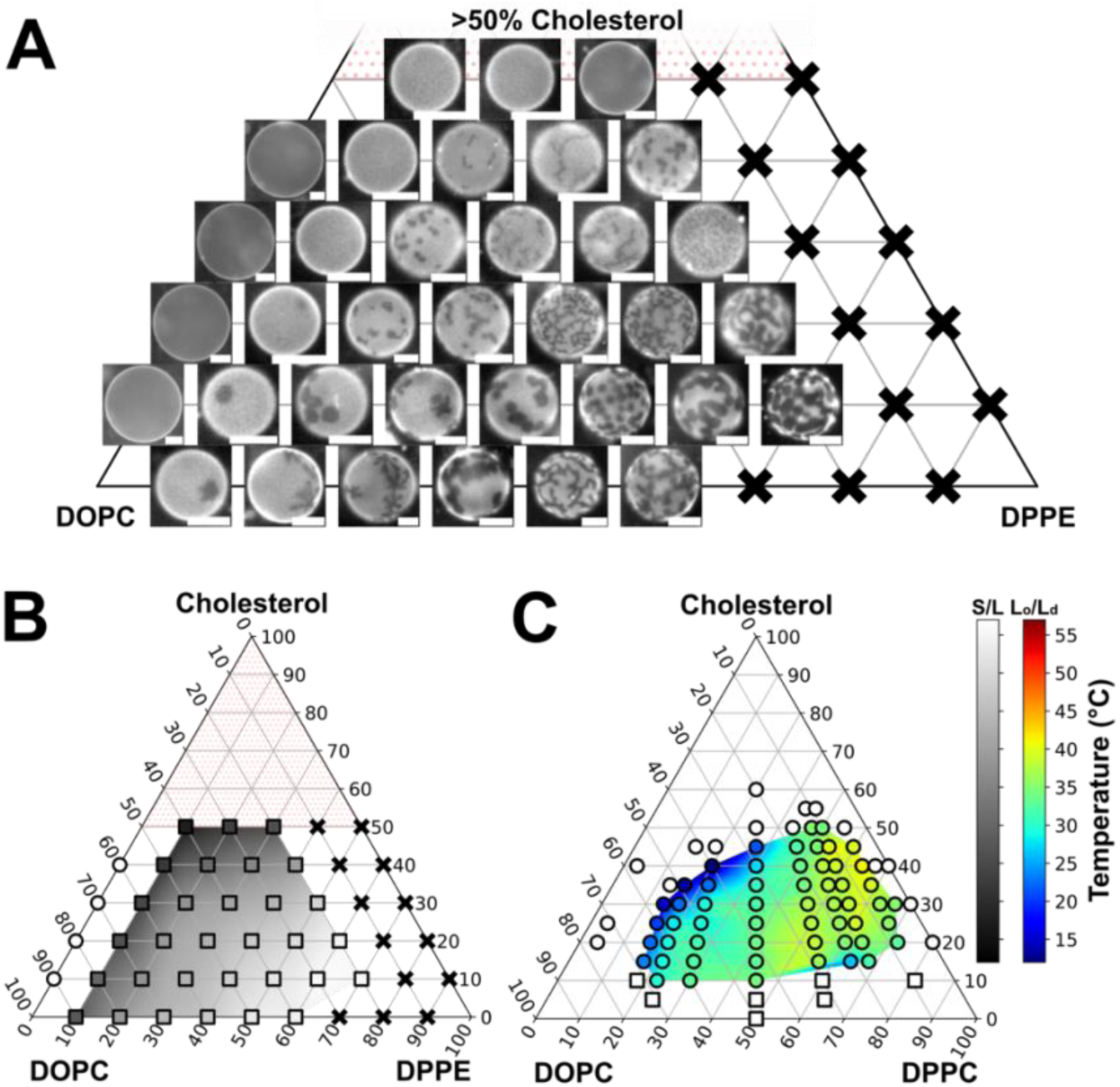
Vesicles of DOPC/DPPE/cholesterol exhibit coexisting solid and liquid phases (rather than coexisting liquid phases). **A)** Representative images at 30°C of vesicles of DOPC/DPPE/cholesterol. Solid phases are dark, and liquid phases are bright. Scale bars: 20 µm. Black X’s mark compositions for which vesicle yields were too small to measure transition temperatures. Cholesterol mole fractions above 50% were avoided (stipple) because they likely exceed maximum cholesterol levels that incorporate in membranes with PE-lipids (49). Wider fields of view with larger images are in Fig. S4-S6. **B)** Transition temperatures in vesicles of DOPC/DPPE/cholesterol, from one liquid phase at high temperature to coexisting solid and liquid phases at lower temperatures (filled squares). Unfilled circles indicate that only one liquid phase was observed. Errors in *T*_mix_ are shown in Fig. S3. Sigmoidal curves used to calculate *T*_mix_ are in Fig. S13-S15. All values are recorded in Table S1. **C)** Transition temperatures in vesicles of DOPC/DPPC/cholesterol, from one liquid phase at high temperature to either coexisting L_o_ and L_d_ phases (filled circles) or coexisting solid and liquid phases (squares), reproduced from (34).

We hypothesized that replacing the low-*T*_melt_ lipid of DOPC with DiphyPC (also known as di(16:0-4Me)PC) might produce liquid-liquid phase separation in membranes containing DPPE (Fig. 4). The methylated chains of DiphyPC give it a very low *T*_melt_ of <-120°C (41). Lipids with isopranyl chains as in DiPhyPC are found in archaebacteria and may allow some organisms to survive in extreme environments (57). Even though DiphyPC is typically associated with robust liquid-liquid phase separation over broad composition ranges of PC-lipids (6, 35), in vesicles containing DPPE, coexisting liquid phases appear over only a small island of mole ratios near the center of the phase diagram (Fig. 4B). Liquid phases are characterized by round domains (Fig. 2C). In taut vesicles, these domains diffuse, collide, and coalesce until only one domain of each phase remains. The liquid ordered (L_o_) phase is identified by being enriched in high-*T*_melt_ lipids and by excluding the dye relative to the liquid disordered (L_d_) phase, as in Fig. S8. When coexisting solid and liquid phases are observed, the area fraction of solid phase increases with DPPE mole fraction, indicating the solid is enriched in DPPE (Fig. 4A, and large fields of view in Fig. S9-S11).

Data along the binary axes in our phase diagrams provide an opportunity to compare with published values. Transition temperatures for data closest to the DPPE-cholesterol axis in Figure 3B (at 20 mol% DOPC) and Figure 4B (at 10 mol% DiPhyPC) are indistinguishable with the onset of solid phases in binary membranes of DPPE-cholesterol measured by Blume and Griffin (who report errors in that liquidus line of at least ±3°C) (52). Similarly, our data for solid-liquid transitions in binary membranes of DOPC and DPPE in Figure 3B are indistinguishable from data by Sakuma et al. (51). In turn, these two data sets are largely in agreement with data by Wu and McConnell (53), with two subtle discrepancies that may be of interest to experts. First, Wu and McConnell’s transition temperatures are within error for mixtures with 50-60% DPPE but are slightly higher (by ≥5°C) for lower DPPE percentages. Second, Wu and McConnell reported a line of three-phase coexistence (one liquid and two solid) at 50°C. This three-phase line is reasonable if two data points supporting the line (namely, changes in the slope of the high field signal height of the paramagnetic resonance spectrum at ∼50°C for membranes with 80% and 85% DPPE) are favored over data that disagree with the line (namely, changes in the slope at 40-46°C for membranes with 15% DPPE) (53). Our data and data from Sakuma et al. would be consistent with a three-phase line only if the line fell below ∼25°C (51).

**Figure 4.**
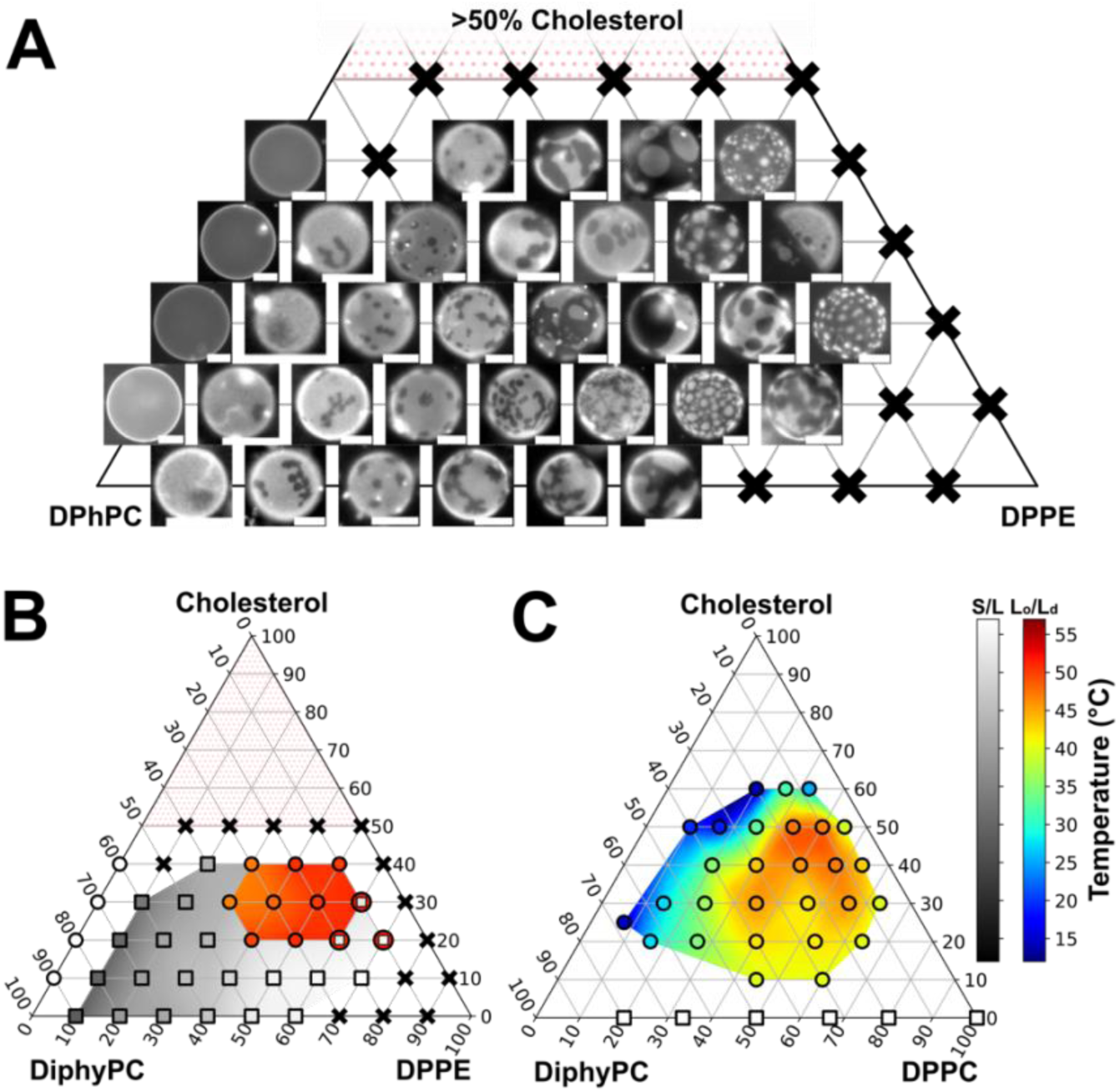
Vesicles of DiPhyPC/DPPE/cholesterol exhibit coexisting liquid phases or coexisting solid and liquid phases. **A)** Representative images at 30°C of vesicles of DiphyPC/DPPE/cholesterol. Dark, noncircular domains are solid. L_d_ domains are bright with respect to both L_o_ and solid domains. Scale bars: 20 µm. Black X’s mark compositions for which vesicle yields were too small to measure transition temperatures. Cholesterol mole fractions above 50% were avoided (stipple) because they likely exceed maximum cholesterol levels that incorporate in membranes with PE-lipids (49). Wider fields of view are in Fig. S9-11. **B)** Transition temperatures in vesicles of DiphyPC/DPPE/cholesterol, from one liquid phase at high temperature to either coexisting solid and liquid phases (grey squares) or coexisting L_o_ and L_d_ phases (red circles). Unfilled circles indicate that only one liquid phase was observed. At three lipid ratios (10:60:30, 10:70:20, and 20:60:20 DiphyPC/DPPE/cholesterol), vesicle membranes were in one liquid phase at high temperature, coexisting solid and liquid phases at an intermediate temperature, and three coexisting phases (solid, L_o_, and L_d_) at a lower temperature. Errors in *T*_mix_ are shown in Fig. S7. Sigmoidal curves used to calculate *T*_mix_ values are in Fig. S16-S18. All values are recorded in Tables S2-S3. **C)** Transition temperatures in vesicles of DiphyPC/DPPC/cholesterol, from one liquid phase at high temperature to either coexisting L_o_ and L_d_ phases (filled circles) or coexisting solid and liquid phases (squares), reproduced from (35). Areas between data points are interpolated and do not reflect uncertainties in data points, which are quoted as midpoints in temperature ranges that span ±2°C (35).

### Replacing DPPC with DPPE makes Lo and solid phases distinguishable by microscopy

An unexpected advantage of incorporating DPPE into ternary membranes is that L_o_ and solid phases become distinguishable by fluorescence microscopy, and subsequent transitions into the 3-phase region (L_o_, L_d_, and solid) are clearly visible (Fig. 5). Previously, 3-phase coexistence has been observed by fluorescence microscopy for only a few combinations of dyes, lipids, and solutions (e.g., (58, 59)). As a result, identification of 3-phase regions by fluorescence microscopy has required observation of changes in domain shapes (60) or in the inability of domains to diffuse and coarsen quickly (35). When L_o_ and solid phases are not distinguishable, alternative spectroscopic techniques (54, 60, 61) or atomic force microscopy (59, 62) must be deployed to conclusively identify the 3-phase region.

Figure 5 shows an example of a liquid phase at a high temperature (57°C) demixing into coexisting liquid and solid phases at a lower temperature (53°C). As temperature decreases still further (48°C), three fluorescence levels are visible, a bright L_d_ phase, a gray L_o_ phase, and a dark solid phase. In many cases, as in Fig. 5, solid domains constrain the motion of the liquid domains. Additional images of vesicles with clear 3-phase coexistence are shown in Fig. S12.

Viewed in the 3-dimensional space of a ternary phase diagram, the volume in which all three phases (L_o_, L_d_, and solid) coexist is most easily imagined as an irregular triangular pyramid, with its tip pointing up toward high temperatures. In traditional ternary membranes with PC-lipids (e.g., DOPC/DPPC/cholesterol), the pyramid is shaped like a shard lying on its side, with two long edges defined by long tie-lines between the L_d_ phase and the other phases (61). In contrast, in the DiphyPC/DPPE/cholesterol phase diagram in Fig. 4B, the pyramid must be more compact because the L_o_/L_d_ coexistence region is smaller. The face of the pyramid that touches the L_o_/solid coexistence region must have a shallow slope because transitions as in Fig. 5 (from a 2-phase, L_o_/solid region to a 3-phase, L_o_/L_d_/solid region) occur over lipid compositions that span at least 10 mol%.

Because L_o_ and solid phases are distinguishable by fluorescence microscopy, we can learn more about how the substitution of DPPE for DPPC results in a phase diagram with primarily solid-liquid coexistence (as in Fig. 4), as opposed to a diagram with only liquid-liquid coexistence (as in DiphyPC/DPPC/cholesterol (35)). The change from DPPE to DPPC involves the addition of only three methyl groups to the lipid headgroup. In Fig. 6, we add these methyl groups one-by-one for three lipid ratios. Addition of each methyl group is expected to decrease the monolayer spontaneous curvature due to the lipid, based on trends for DOPE, DOPE-1Me, and DOPC (63). We find that largest change in the phase behavior occurs upon addition of the first methyl group, from DPPE to singly-methylated DPPE (“1Me”), whereupon temperatures of the 2-phase (solid/liquid) and 3-phase (L_o_/L_d_/solid) transitions become nearly indistinguishable (Fig. 6).

**Figure 5.**
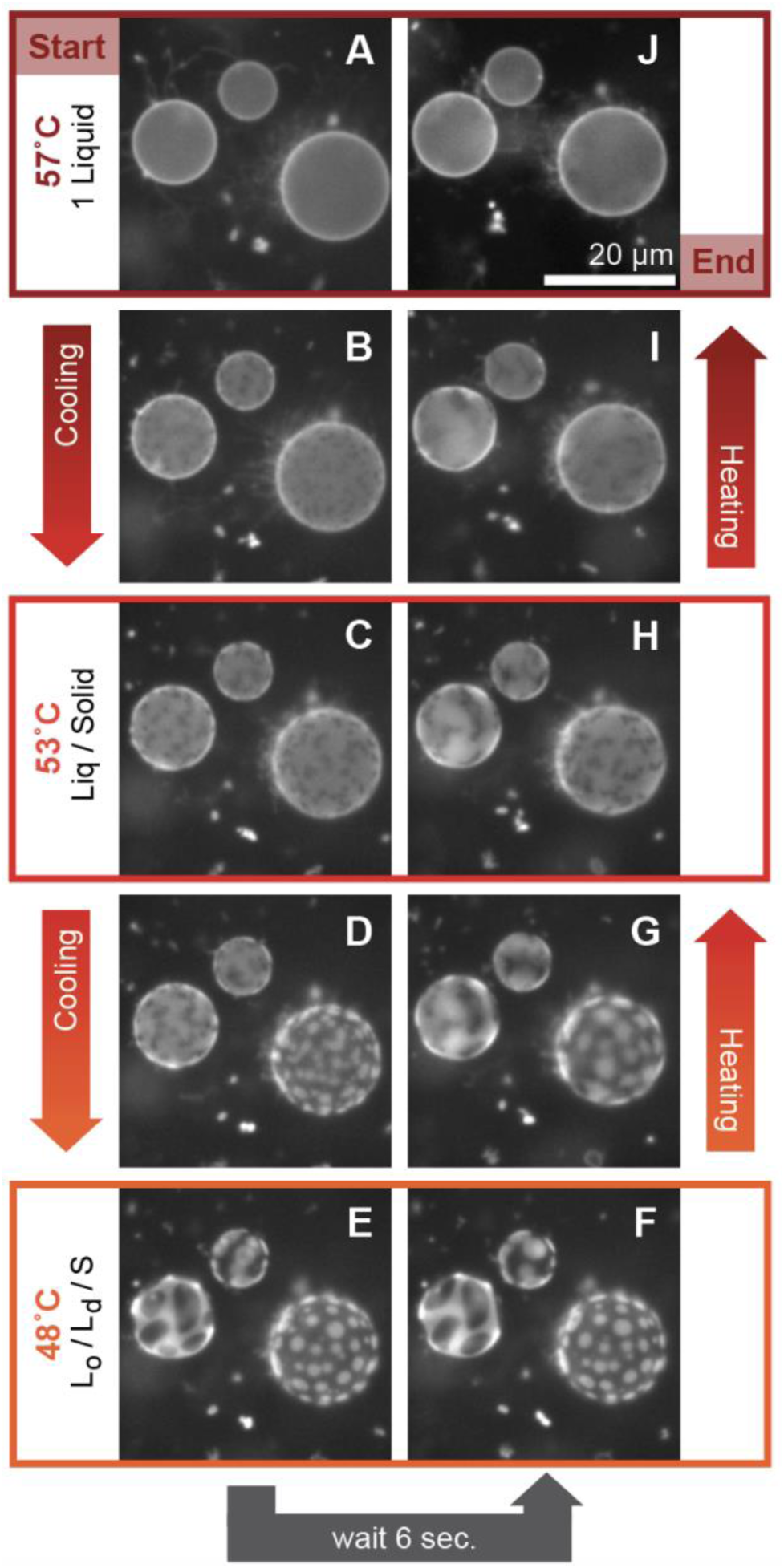
Liquid-liquid phase separation occurs at lower temperature than solid-liquid separation in 10:70:20 DiPhyPC/DPPE/cholesterol membranes. A) Vesicle membranes are in one liquid phase at high temperature. B) Solid domains nucleate at ∼56°C. C) Domains of dark, noncircular, solid phase coexist with a bright, liquid phase. D) At ∼52°C, the liquid phase begins demixing into L_o_ and L_d_ phases. The L_o_ phase is identified by an intermediate fluorescence level between the dark solid phase and bright L_d_ phase. E) At 48°C, micron-scale regions of three fluorescence levels are observed, corresponding to solid, L_o_, and L_d_ phases. Solid phases are wet by L_o_ domains. E-F) With time, domains diffuse, and liquid domains can merge. F-J) Phase transitions are reversible with temperature. Images correspond to Video S1; timepoints for images A–J are 0, 16, 31, 40, 48, 54, 70, 76, 96 and 119 sec, respectively. These transitions occur in the same order in vesicles of 10:60:30 and 20:60:20 DiphyPC/DPPE/cholesterol.

**Figure 6.**
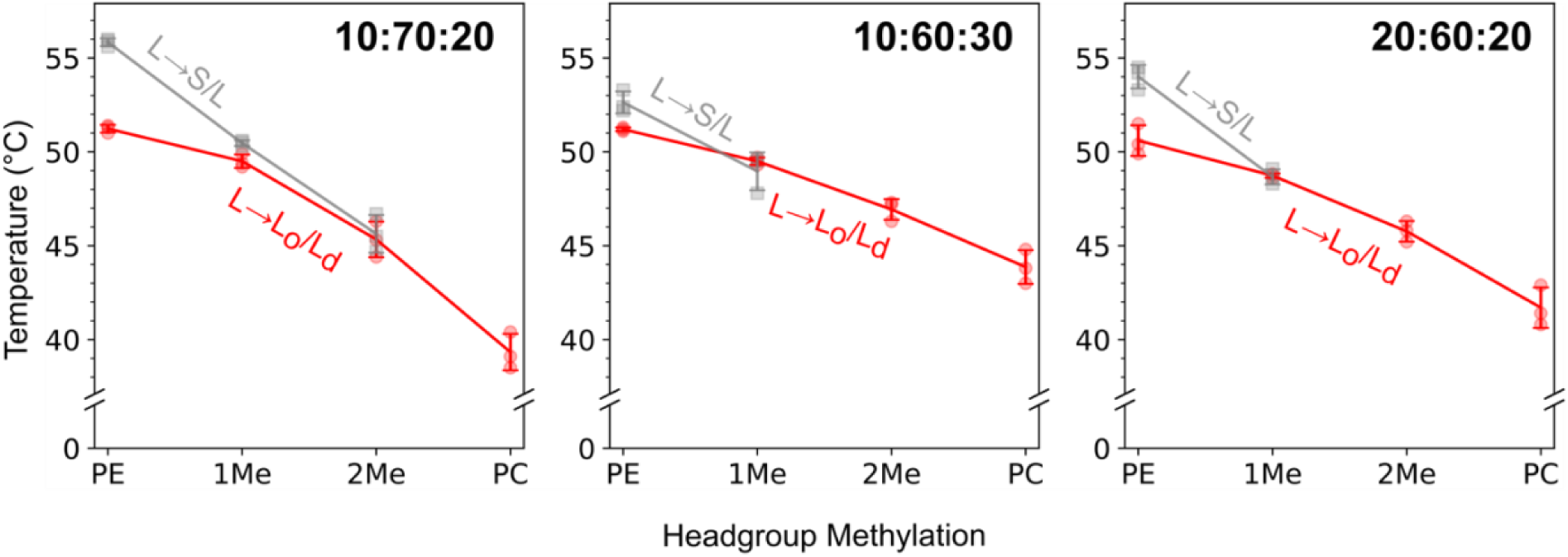
Methylation of DPPE headgroup eradicates solid-liquid coexistence at high temperatures. As temperature is decreased in vesicles of 10:70:20, 10:60:30, and 20:60:20 DiphyPC/DPPE/cholesterol (denoted “PE” on each x-axis) transitions are observed from one liquid phase at high temperature, to coexisting solid and liquid phases, and then to three phases (L_o_, L_d_, and solid). In each of the three mixtures, DPPE was replaced by singly-(“1Me”), doubly-(“2Me”), or triply-methylated (“PC”) versions of the molecule. In all three panels, gray squares show temperatures at which coexisting solid and liquid phases appear, and red circles show temperatures at which both L_o_ and L_d_ phases appear, for three independent measurements. Error bars are standard deviations of the triplicate measurements. Lines through triplicate averages are to guide the eye and are not fits to a model. Corresponding data are tabulated in Tables S2-8.

### Why do membranes with DPPE exhibit so little liquid-liquid phase separation?

The lack of liquid-liquid phase separation in Fig. 3A-B (and the limited range of it in Fig. 4A-B) highlight how difficult it is to predict ternary phase diagrams based on physical attributes of single lipid types or even from binary phase diagrams. This challenge exists even when all phospholipids have PC-headgroups. For example, for the system of DiPhyPC/DPPC/ cholesterol, none of the binary membranes have large-scale, coexisting phases above 42°C (34, 51), the melting temperature of DPPC (10). An intuitive, but incorrect, conclusion would be that all ternary mixtures between these binary axes should exhibit one liquid phase. Instead, an extensive, closed-loop region of liquid-liquid coexistence appears above 42°C (35). Similarly unintuitive behavior occurs below 42°C. Binary mixtures of DiPhyPC/DPPC produce solid phase boundaries (liquidus lines) indistinguishable from those of DOPC/DPPC (within experimental error, for DPPC ≥ 20 mol%) (51, 64). Nevertheless, liquid-liquid coexistence is observed in ternary membranes of DiPhyPC/DPPC/cholesterol over wider ranges of compositions at all temperatures than in DOPC/DPPC/cholesterol.

Likewise, the lack of liquid-liquid phase separation in Fig. 3A is surprising because DPPE’s melting temperature (*T*_melt_ = 62.3 ± 5.0°C (22)) is close to the melting temperature of DAPC (di(20:0)PC, *T*_melt_ = 65.3 ± 1.5°C (10)), and coexisting liquid phases persist in equivalent ternary membranes with DAPC (34). Increasing the *T*_melt_ of PC-lipids or sphingomyelins in ternary mixtures tends to simply increase liquid-liquid transition temperatures (11, 34, 60) rather than suppress liquid-liquid coexistence. Instead, Fig. 3A is reminiscent of ternary phase diagrams in which palmitoyl-ceramide (PCer) is the high melting temperature lipid (14). This is surprising because PCer is structurally very different from DPPE, namely, PCer has no polar headgroup at all, which gives it an extraordinarily high melting temperature of 90.0°C (65). Unexpected phase behavior, whether it is observed in membranes with PC-lipids or with PE-lipids, highlights the importance of experimental measurement of membranes with new lipid combinations.

One clear difference between DPPE and DPPC is in the monolayer spontaneous curvatures they impart to membranes. PE-lipids and cholesterol are both cone-shaped (66). Several authors have discussed how cone-shaped lipids could give rise to sub-micron heterogeneities in bilayers with excess area (e.g. (67, 68)). The effect of lipid spontaneous curvature on large-scale phase separation may depend on whether the PE-lipid predominantly resides in the L_d_ phase or in the L_o_ phase. In some membranes, including symmetric membranes, small-scale heterogeneity may be required for large-scale L_o_ phases. In all atom-simulations of symmetric bilayers, lipid sub-structures were observed in the L_o_ phase (69). For membranes in which different lipid types have similar spontaneous curvatures (as for PE-lipids and cholesterol), the lipids would not contribute to L_o_ sub-structures, whereas lipids with dissimilar spontaneous curvatures (as for PC-lipids and cholesterol) would (67, 70). If sub-structures are necessary for L_o_ phases, a symmetric membrane enriched in DPPE would be less likely to demix into coexisting liquid phases, consistent with Fig. 3B.

Differences in lipid spontaneous curvature between PE-lipids and PC-lipids may arise from differences in headgroup structural volumes and differences in hydrogen bonding. Whereas PC-lipids in bilayers hydrogen-bond only with surrounding water molecules, PE-lipids also hydrogen bond with amine protons of neighboring PE headgroups, observed through a C=O stretching band (71). These differences in structure and hydrogen bonding are cited as reasons that PE-lipids have the lowest molecular areas and lowest calculated hydrations of all glycerophospholipids (72).

At first glance, the data in Figure 6 imply that solid phases persist at high temperatures in membranes of DPPE because of hydrogen bonding between molecules, which is missing in singly-(1Me), doubly-(2Me), or triply-(PC) methylated lipids. However, a solid phase persists to ∼1°C above coexisting liquid phases in membranes of 10:70:20 DiphyPC/1Me/cholesterol (Fig. 6). Disambiguating effects due to headgroup size and hydrogen-bonding is difficult because both parameters change when DPPE is replaced by a singly-methylated lipid (1Me). If hydrogen bonding were the most important variable to consider, then we would expect spontaneous monolayer curvatures of singly-(1Me) methylate lipids to be closer to those of DOPC than of DOPE, which is not supported by data (23). Lewis and McElhaney’s caveat that

“Although many of the physical differences between hydrated PC and PE bilayers are attributed to… the capacity of the headgroup amine group of PE to hydrogen bond with other acceptor groups on the lipid molecule… to our knowledge there is little direct experimental support for such a premise” feels as relevant today as when they wrote it decades ago (71).

### Outward tubules are common

Electroforming ternary vesicles with DPPE presents technical challenges. As noted earlier, vesicles with high fractions of DPPE have low yields. In addition, most fields of view have at least one high-curvature tubule and/or a bright aggregate of lipids (Fig. S4-S6 and S8-S11). A worst-case scenario is shown in Fig. 2D. Although tubules and aggregates can appear in corresponding samples with PC-lipids, they are less common (34, 35). Imperfections of tubules and aggregates do not appear to significantly affect transition temperatures or domain morphology. As noted earlier, there is good agreement between the onset of solid phases in binary DOPC-DPPE electroformed vesicles (Fig. 3B) and in equivalent vesicles formed by gentle hydration, for which tubules were not reported (51). In Fig. 5, the tubules seem unchanged upon cooling (panels A-B) or heating (panels I-J) through a liquid-liquid phase transition. As expected, an increase in the fraction of the surface area covered with solid and Lo phases causes many tubules appear to be incorporated into vesicles (panel E), and the process is reversible (73).

Although tubules are not a focus of this study, they are perplexing. Tubulation is known to be enhanced by temperature fluctuations near solid-liquid and liquid-liquid transitions as can happen when a temperature controller’s set point is near the transition (73, 74) and by inclusion of cholesterol, which can reduce the temperature of the lamellar-H_II_ transition by 20°C (18). As seen in other systems, partitioning of the *L*_d_-probe into tubules implies they are in the *L*_d_ phase (75), which in turn implies the tubules are depleted of DPPE. Nearly all tubules in Fig. 2 point outward, unlike tubules observed by Sakuma et al., which point inward for binary DMPE/DPPC vesicles formed by gentle hydration (73). An outward orientation implies a compositional asymmetry with more PE-lipids in the inner leaflet. This is opposite the direction expected from Steinkühler et al., who found evidence that electroformation drives negatively charged lipids to the outer leaflet of vesicles (76). Although “PE bears little net charge at neutral pH”, its pKa of 9.6 ± 0.1 means a (small) fraction of PE-lipids should be negatively charged (77). Any asymmetry in lipid composition should be at least partially mitigated by the high temperature (75°C) at which vesicles were electroformed (76).”

## CONCLUSIONS

In this manuscript, we expanded the known set of phase diagrams of ternary membranes to include membranes with PE-lipids. Specifically, we replaced a PC-lipid that is common in ternary diagrams (DPPC) with its corresponding PE-lipid (DPPE). Changes to the molecular structure are small: these two lipids differ by only three methyl groups. However, subsequent changes to miscibility phase diagrams of ternary membranes containing these lipids are large. Whereas liquid-liquid phase separation is present in vesicles across a wide band of compositions composed of DOPC/DPPC/cholesterol, it is entirely absent from the phase diagram of vesicles of DOPC/DPPE/cholesterol, and only a small island of liquid-liquid phase separation appears in the phase diagram of vesicles of DiphyPC/DPPE/cholesterol. Inside that island, membranes of a small set of compositions begin first in a liquid phase at high temperature, then separate into coexisting solid and liquid phases as temperature decreases, and then undergo a transition into a three-phase region after temperature decreases further. This three-phase region contains coexisting liquid-ordered, liquid-disordered, and solid phases, all of which are distinguishable by fluorescence microscopy. In most other types of lipid membranes, transitions into the three-phase region are difficult to observe by microscopy, due to solid phases nucleating within liquid-ordered phases and/or due to poor contrast of fluorophores between liquid-ordered and solid phases. Ideally, our investigation of ternary membranes containing DPPE will serve as a starting point for other researchers to probe liquid-liquid phase separation in membranes containing other PE-lipids beyond DPPE that are relevant and interesting. Of high interest would be PE-lipids with unsaturated chains and with vinyl linkages to the chains (plasmalogens), both of which are prevalent in biological membranes and have been the subject of molecular dynamics simulations (3, 4, 33, 78–81).

## AUTHOR CONTRIBUTIONS

GJG, CEC, and SLK designed research and protocols. GJG, CLAC, and SN performed research. GJG, SN, and AMM analyzed data. GJG and SLK wrote the manuscript. GJG and SLK edited the manuscript.

## ACKNOWLEDGEMENTS

This research was supported by National Science Foundation grant MCB-1925731 and MCB-2325819 to SLK. The authors thank Bob Weng for preliminary experiments. We thank our anonymous reviewers for constructive feedback, and we thank Alex Sodt, Sergei Sukharev, and Mark Uline for discussions about spontaneous curvature, hydrogen bonding, and phenomenological models, respectively.

## Supporting Material

**Figure S1.**
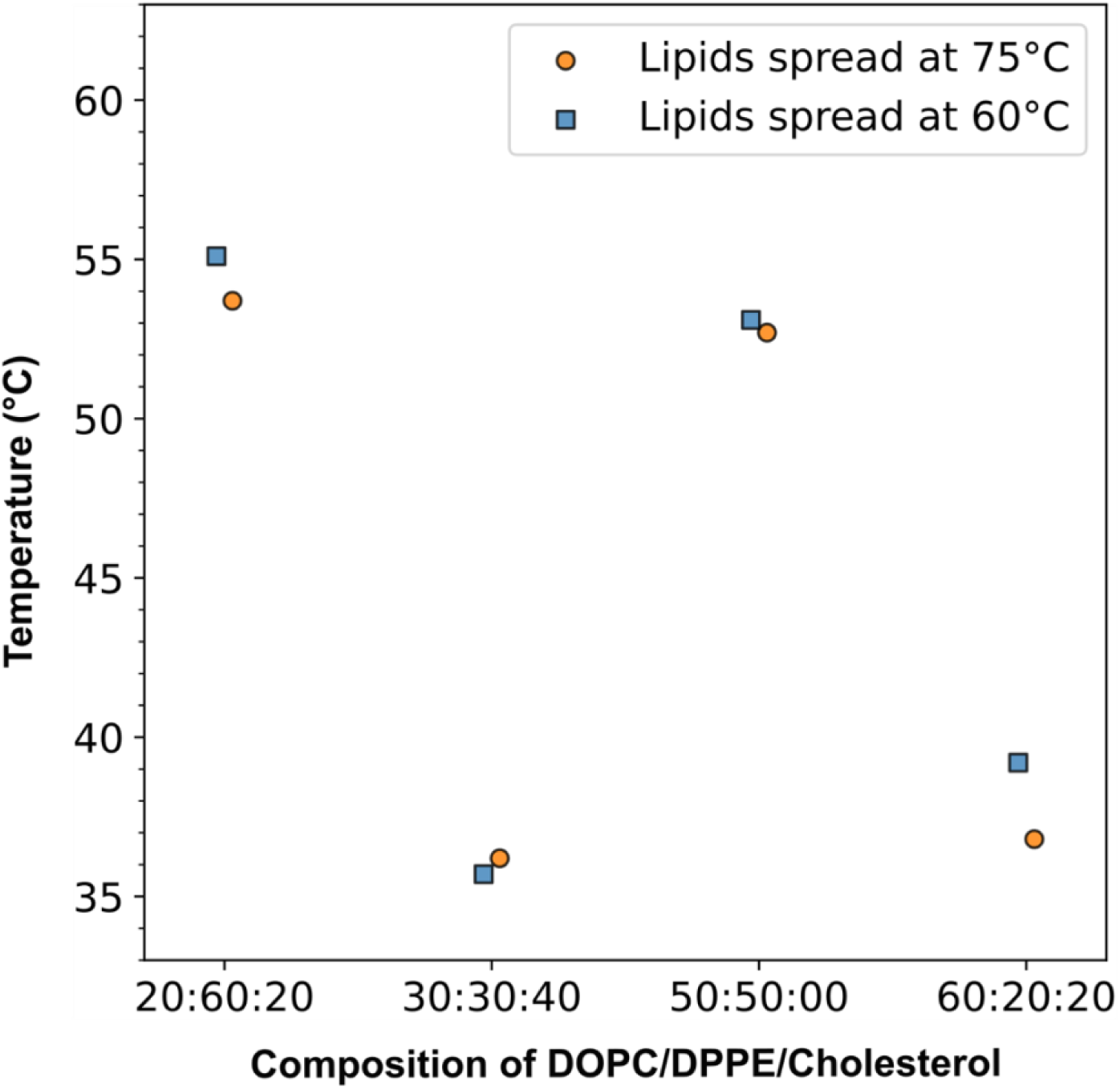
Lipid films spread on slides at 60°C or 75°C (and then electroformed at 75°C) produce vesicles with similar *T*_mix_ values. In all experiments in the main text, vesicles were electroformed at 75°C. This temperature was chosen to be well above the *T*_melt_ of DPPE, which is 62.3 ± 5.0°C (22). Ideally, to prepare lipid films for electroformation, they should have been spread on indium-tin-oxide slides at the same high temperature. However, it is technically difficult to do so, given that chloroform boils at 61°C. Therefore, we assessed if spreading the films at 60°C, a temperature just below chloroform’s boiling point, affected the miscibility phase transition temperature of subsequently electroformed vesicles. We would expect bigger effects for mixtures with higher mole fractions of DPPE, which are the first three mixtures in the plot. For these mixtures, differences between *T*_mix_ of vesicles made from films spread at 60°C and 75°C are inconsequential, and justify our decision to spread all films at 60°C. We were perplexed but not concerned by the larger difference in *T*_mix_ for the mixture with 20% DPPE (the rightmost mixture in the plot). For all four samples, the average difference in *T*_melt_ between samples spread at 60°C and 75°C is ±1.2°C (range 0.4°C to 2.4°C). For comparison, sample-to-sample variations are ±0.1 to ±1.3°C (range 0.1°C to 2.8°C).

**Figure S2.**
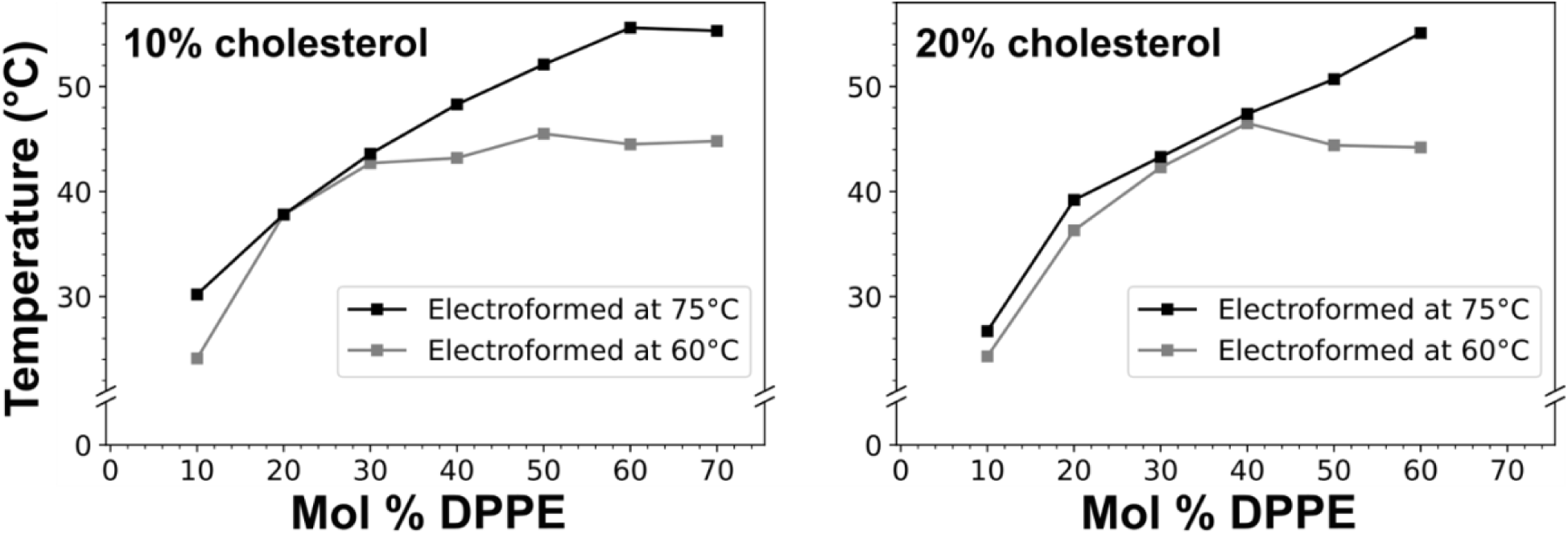
DOPC/DPPE/cholesterol vesicles electroformed at 60°C (below the *T*_melt_ for DPPE) exhibit lower *T*_mix_ values than vesicles electroformed at 75°C, especially at higher DPPE mole fractions. In all experiments in the main text, vesicles were electroformed at 75°C, well above *T*_melt_ for DPPE (62.3 ± 5.0°C (22)). To illustrate the importance electroforming above *T*_melt_, several ternary lipid mixtures composed of DOPC/DPPE/cholesterol were electroformed at 60°C, a temperature commonly used for electroforming vesicles containing PC-lipids such as POPC, DOPC, and DPPC. At higher DPPE compositions, the measured Tmix value is as much as ∼10°C lower for samples electroformed at 60°C than at 75°C. These results imply that insufficient DPPE is incorporated into vesicles at 60°C and are consistent with (43).

**Figure S3.**
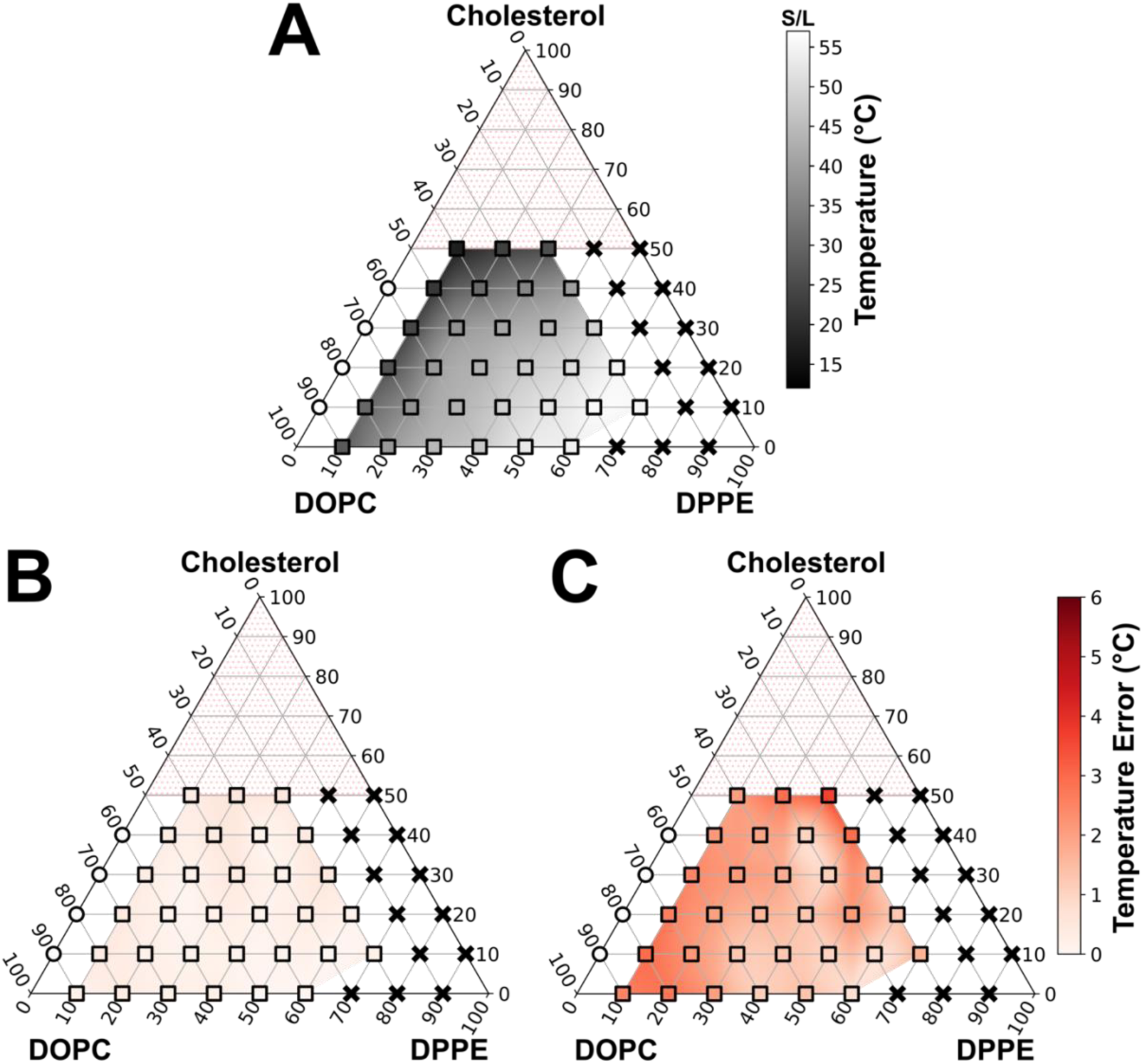
Two methods of assessing error from single measurements of *T*_mix_ for vesicles of DOPC/DPPE/Cholesterol. **A)** Temperatures of phase transitions from one liquid at high temperature to coexisting solid and liquid phases at lower temperatures, from Fig. 3B. **B)** Error in the *T*_mix_ fit parameter calculated from 95% confidence intervals of the sigmoidal fits (Fig. S13-S15), centered at *T* = *T*_mix_. This method yields very small errors in *T*_mix_ (<0.5°C), indicating the sigmoidal fits are good. However, the errors do not give information about the breadth of the transition. **C)** Error in *T*_mix_ calculated as ½(|*T*_90%_ *– T*_10%_|), which is half the difference between the temperature at which 90% of vesicles are phase separated and that at which 10% are phase separated. Errors calculated by this method give more information about sample preparation and the shape of the phase boundary. Smaller errors reflect more uniformity in vesicle-to-vesicle composition and/or more constant transition temperatures from composition to composition. In the absence of independent measurements to measure standard deviations, this method can provide a useful way to assess experimental error. Corresponding data are tabulated in Table S1.

**Figure S4.**
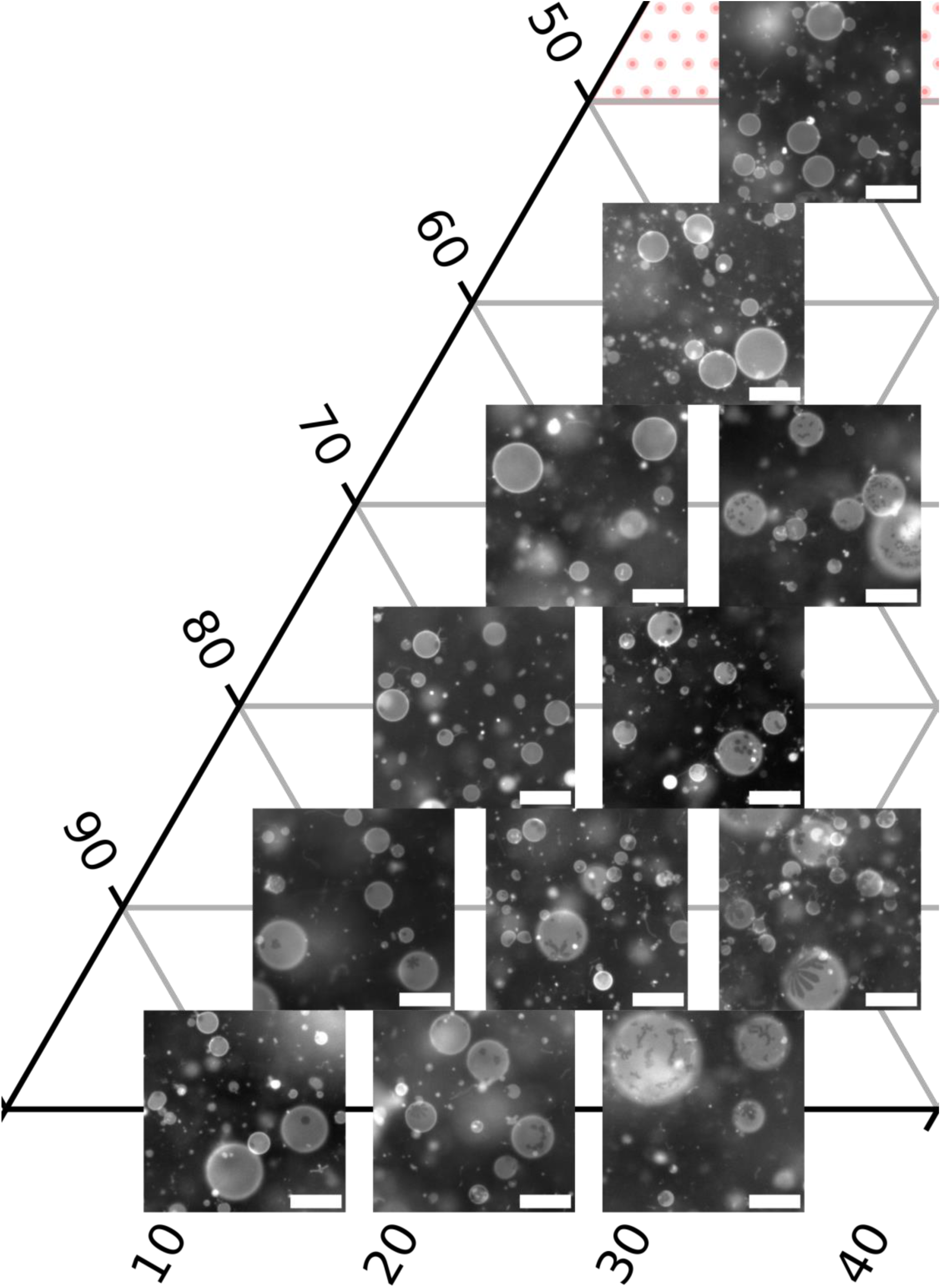
Representative fields of view for vesicles of DOPC/DPPE/cholesterol, covering the left third of the phase diagram in Figure 3b of the main text. The vertex at the left represents 100% DOPC, and high cholesterol is toward the top. Scalebars are 50 µm.

**Figure S5.**
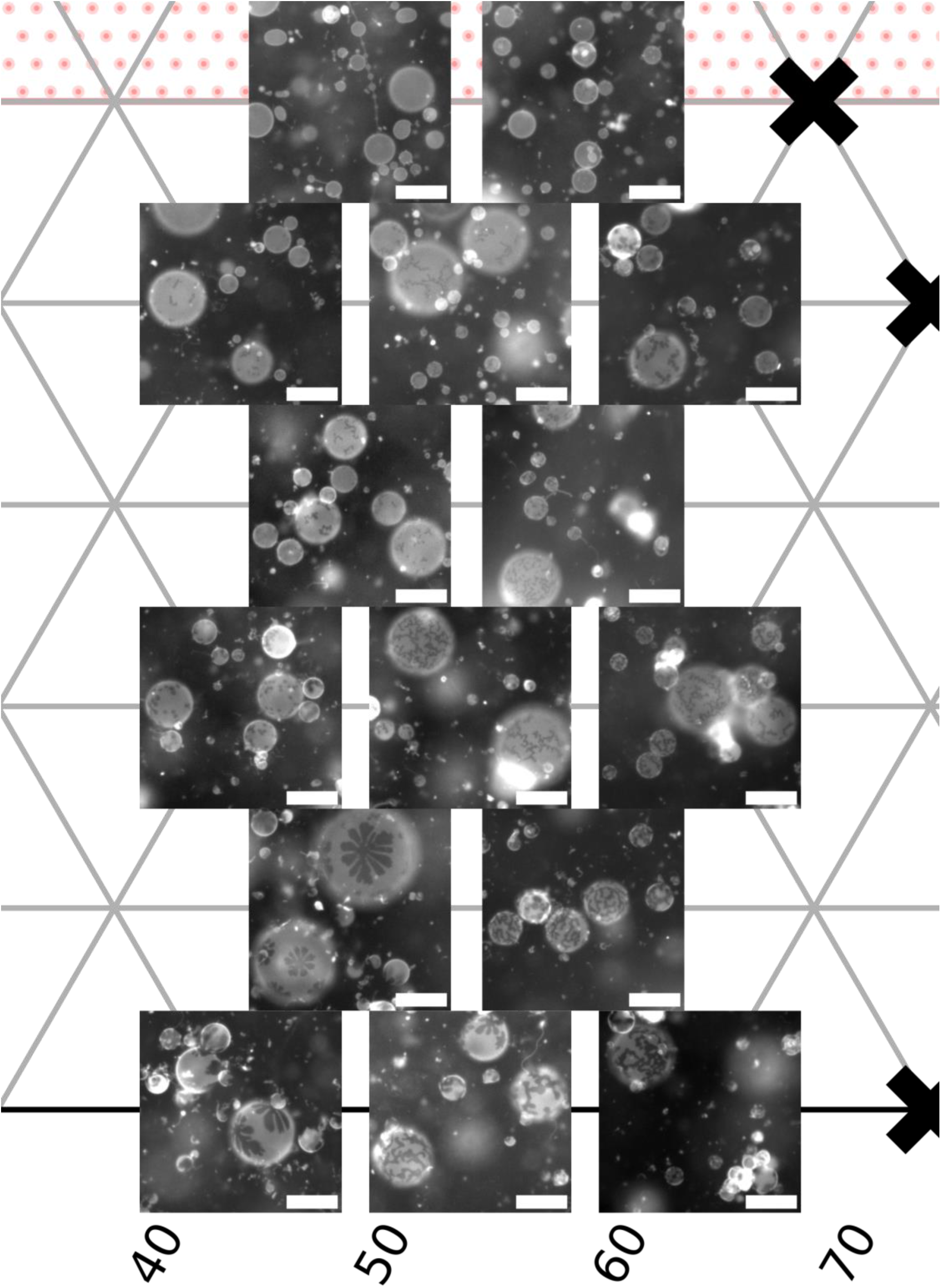
Representative fields of view for vesicles of DOPC/DPPE/cholesterol, covering the middle third of the phase diagram in Figure 3b of the main text. High DOPC fractions are on the left, high DPPE fractions on the right, and high cholesterol toward the top. Scalebars are 50 µm.

**Figure S6.**
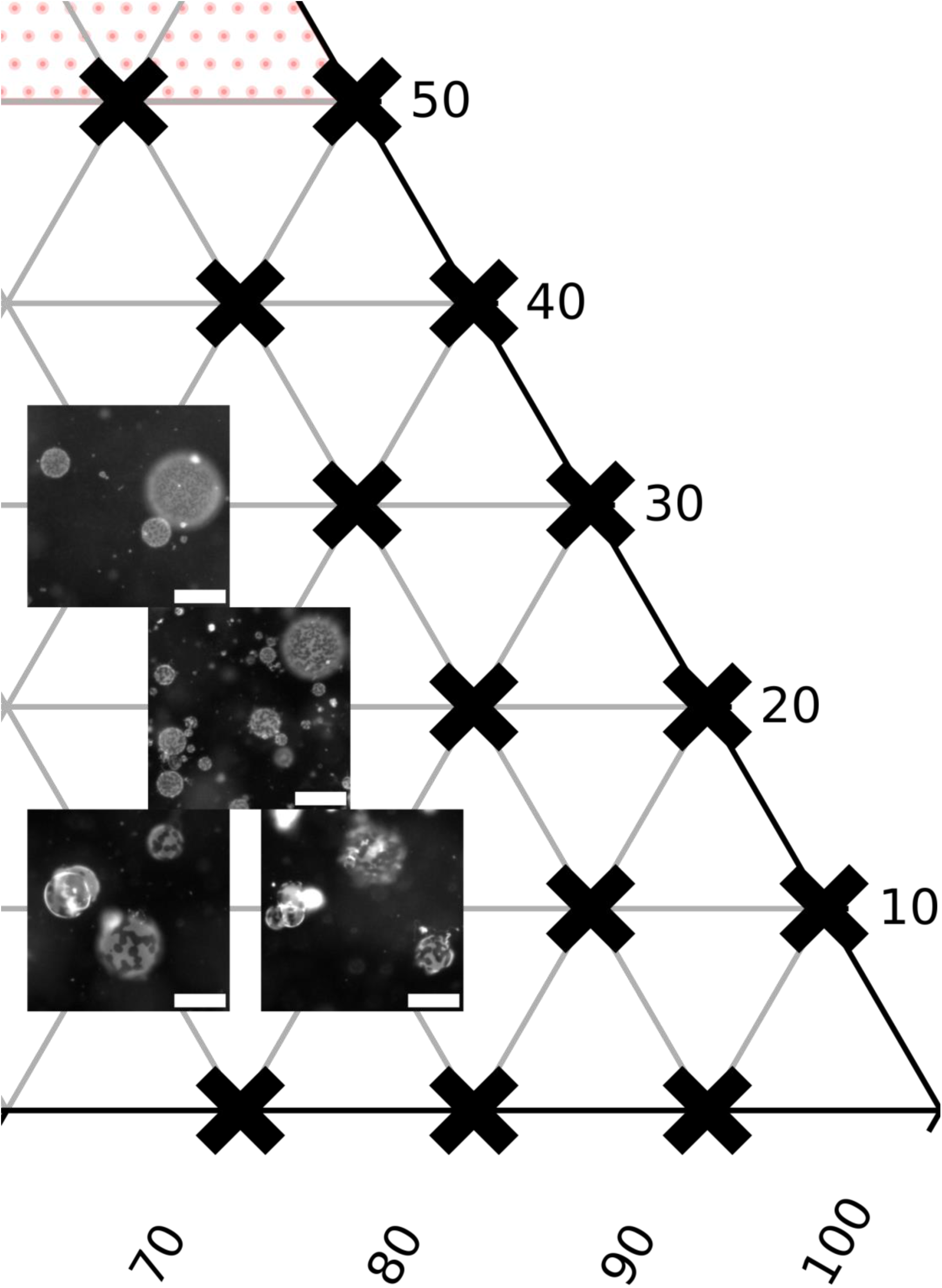
Representative fields of view for vesicles of DOPC/DPPE/cholesterol, covering the right third of the phase diagram in Figure 3b of the main text. The vertex at the right represents 100% DPPE, and high cholesterol is toward the top. Scalebars are 50 µm.

**Figure S7.**
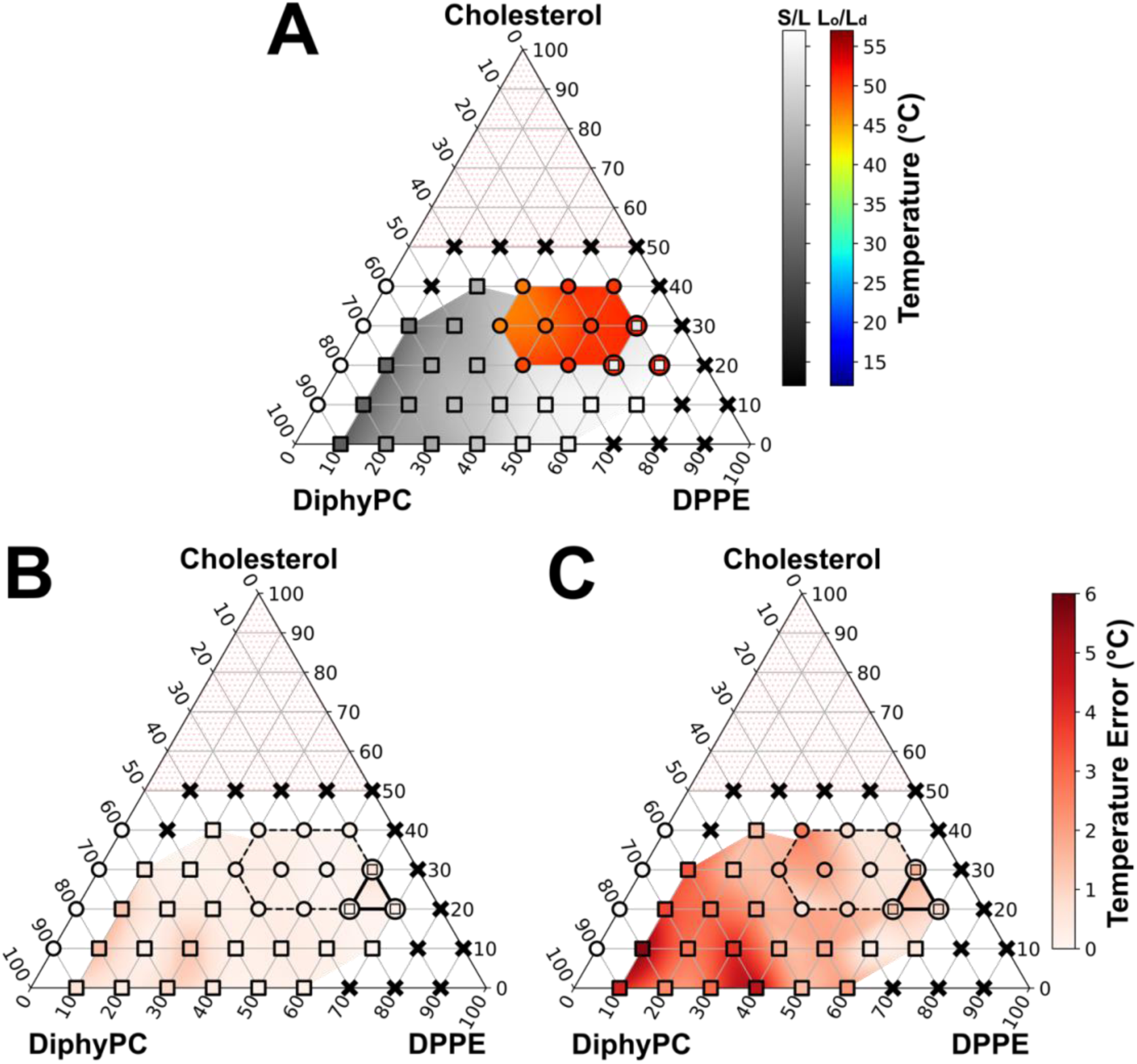
Two methods of assessing error from single measurements of *T*_mix_ for vesicles of DiphyPC/DPPE/Cholesterol. **A)** Temperatures of phase transitions from one liquid to coexisting solid and liquid phases (squares) or to coexisting L_o_ and L_d_ phases (circles), from Fig. 4B. **B)** Error in the *T*_mix_ fit parameter calculated from 95% confidence intervals of the sigmoidal fits (Fig. S16-S18), centered at *T* = *T*_mix_. This method yields very small errors in *T*_mix_ (typically <0.5°C), indicating good sigmoidal fits. However, the errors do not give information about the breadth of the transition. Compositions for which the transition is from one liquid phase to coexisting L_o_ and L_d_ phases are bounded by a dotted black line. Compositions for which we could clearly observe a transition into a 3-phase region are bounded by a solid black line. **C)** Error in *T*_mix_ calculated as ½(|*T*_90%_ *– T*_10%_|), which is half the difference between the temperature at which 90% of vesicles are phase separated and that at which 10% are phase separated. Errors calculated by this method give more information about sample preparation and the shape of the phase boundary. Smaller errors reflect more uniformity in vesicle-to-vesicle composition and/or more constant transition temperatures from composition to composition. In the absence of independent measurements to measure standard deviations, this method can provide a useful way to assess experimental error. Corresponding data are tabulated in Tables S2 and S3.

**Figure S8.**
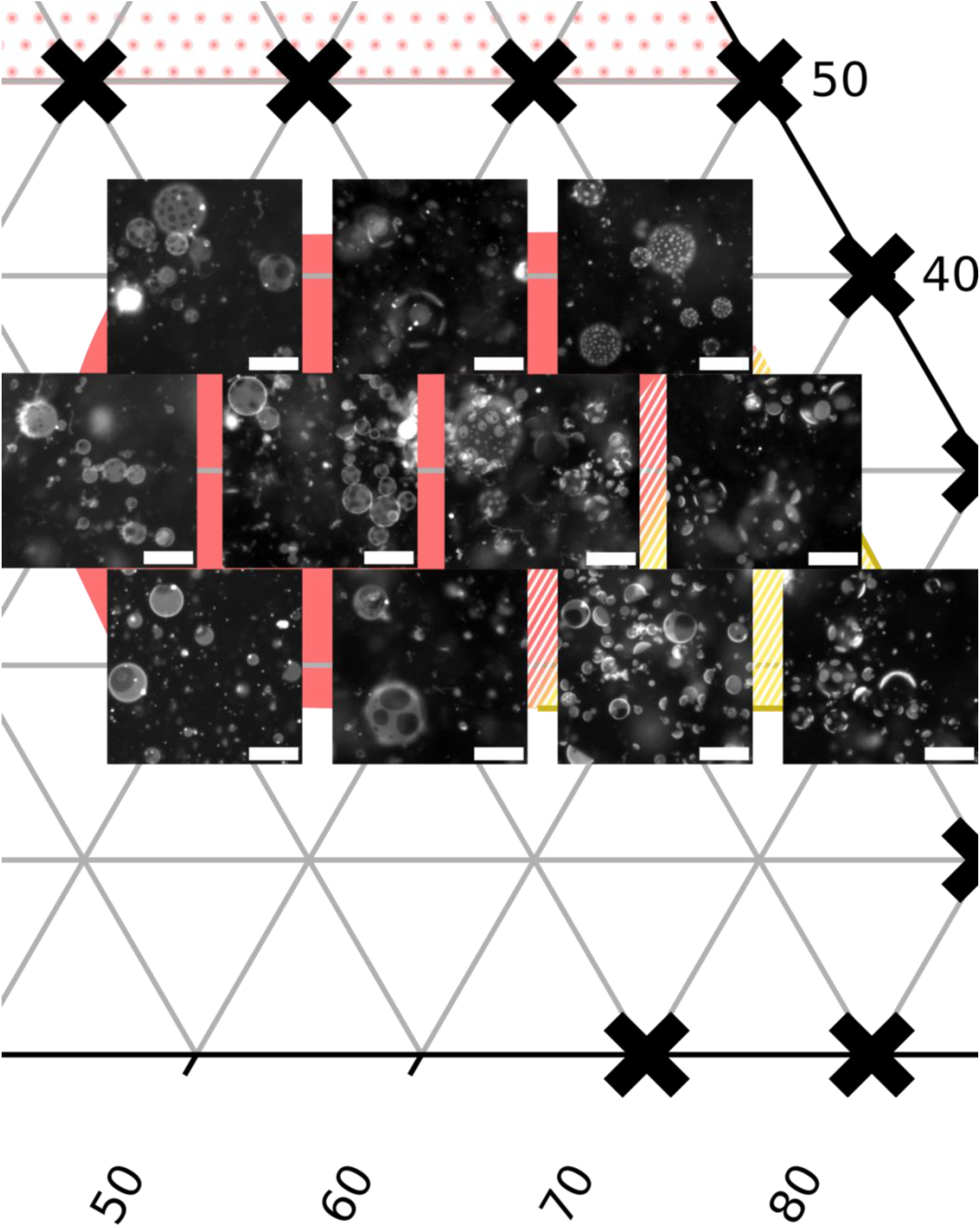
Representative fields of view for vesicles of DiphyPC/DPPE/cholesterol in the liquid-liquid coexistence region in Figure 4b in the main text. Vesicle images were collected between 46-47°C. The region with a red background approximates the L_o_/L_d_ region at this temperature, and the region with a yellow background approximates the 3-phase triangle. Because data were taken in increments of 10 mol%, the fade from red to yellow represents uncertainty in the location of the left edge of the 3-phase triangle. Scalebars are 50 µm.

**Figure S9.**
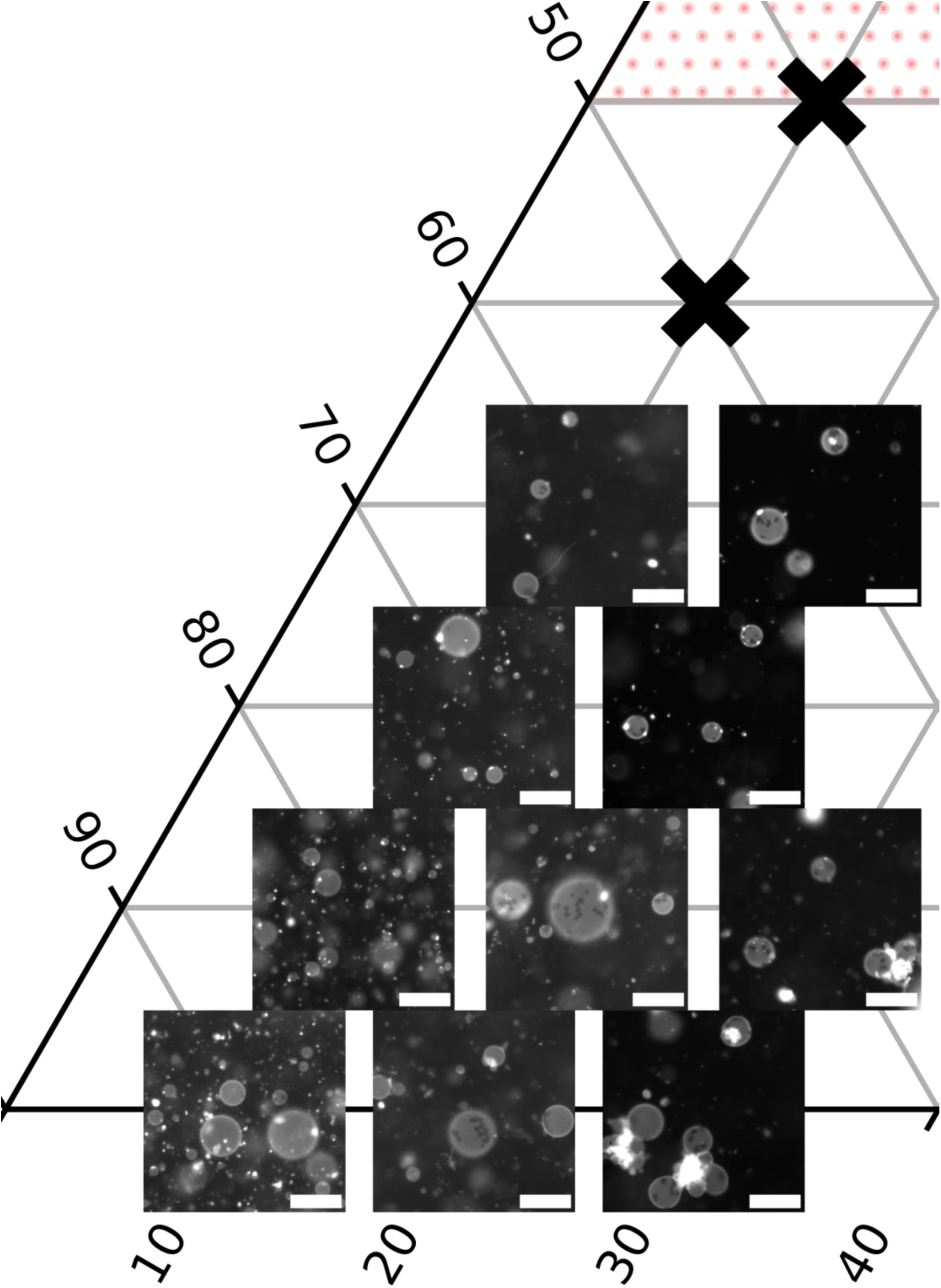
Representative fields of view for vesicles of DiphyPC/DPPE/cholesterol, covering the left third of the phase diagram in Figure 4b of the main text. The vertex at the left represents 100% DiphyPC. High cholesterol is toward the top. Scalebars are 50 µm.

**Figure S10.**
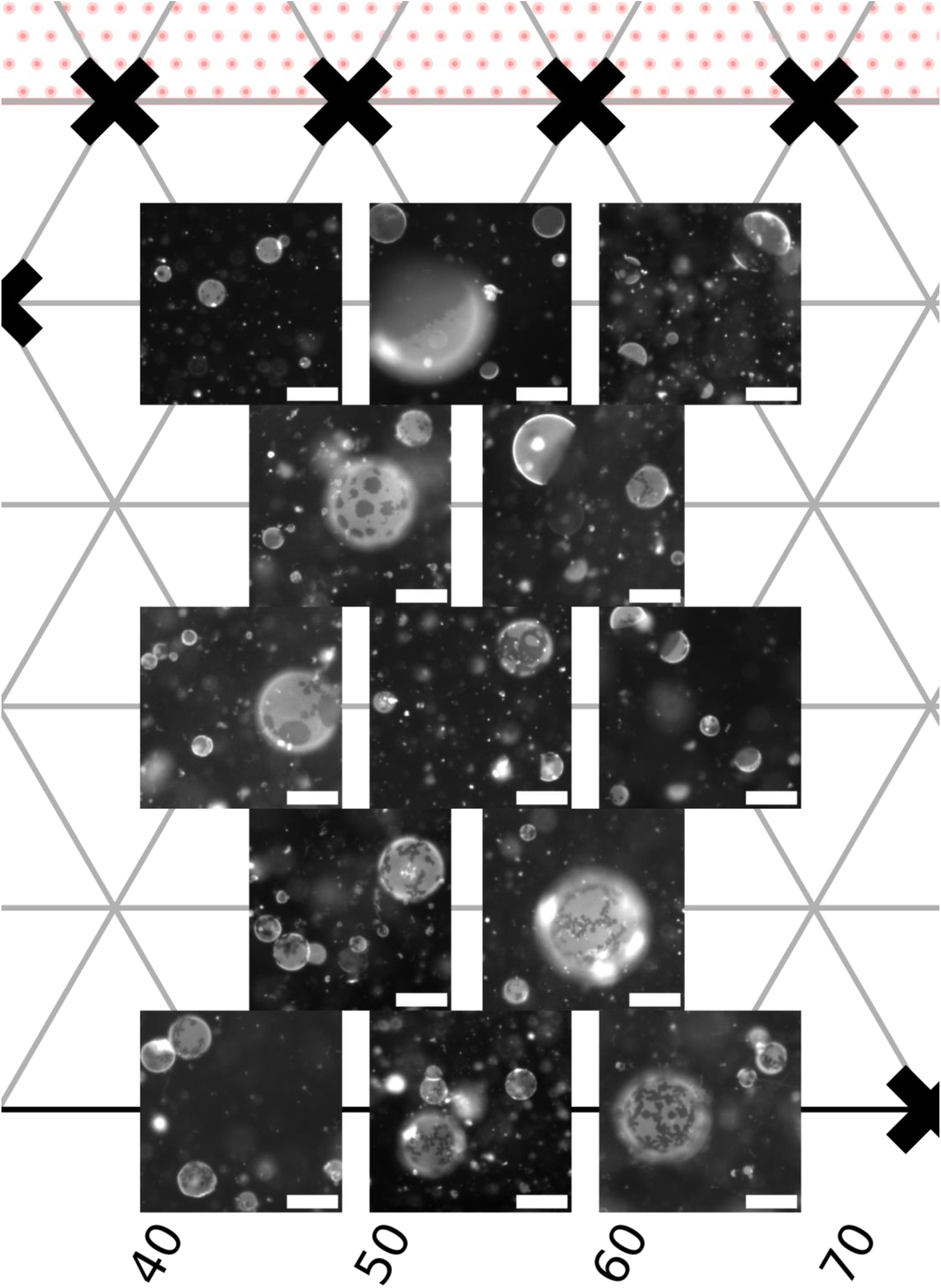
Representative fields of view for vesicles of DiphyPC/DPPE/cholesterol, covering the middle third of the phase diagram in Figure 4b of the main text. High DiphyPC fractions are on the left, high DPPE fractions on the right, and high cholesterol toward the top. Scalebars are 50 µm.

**Figure S11.**
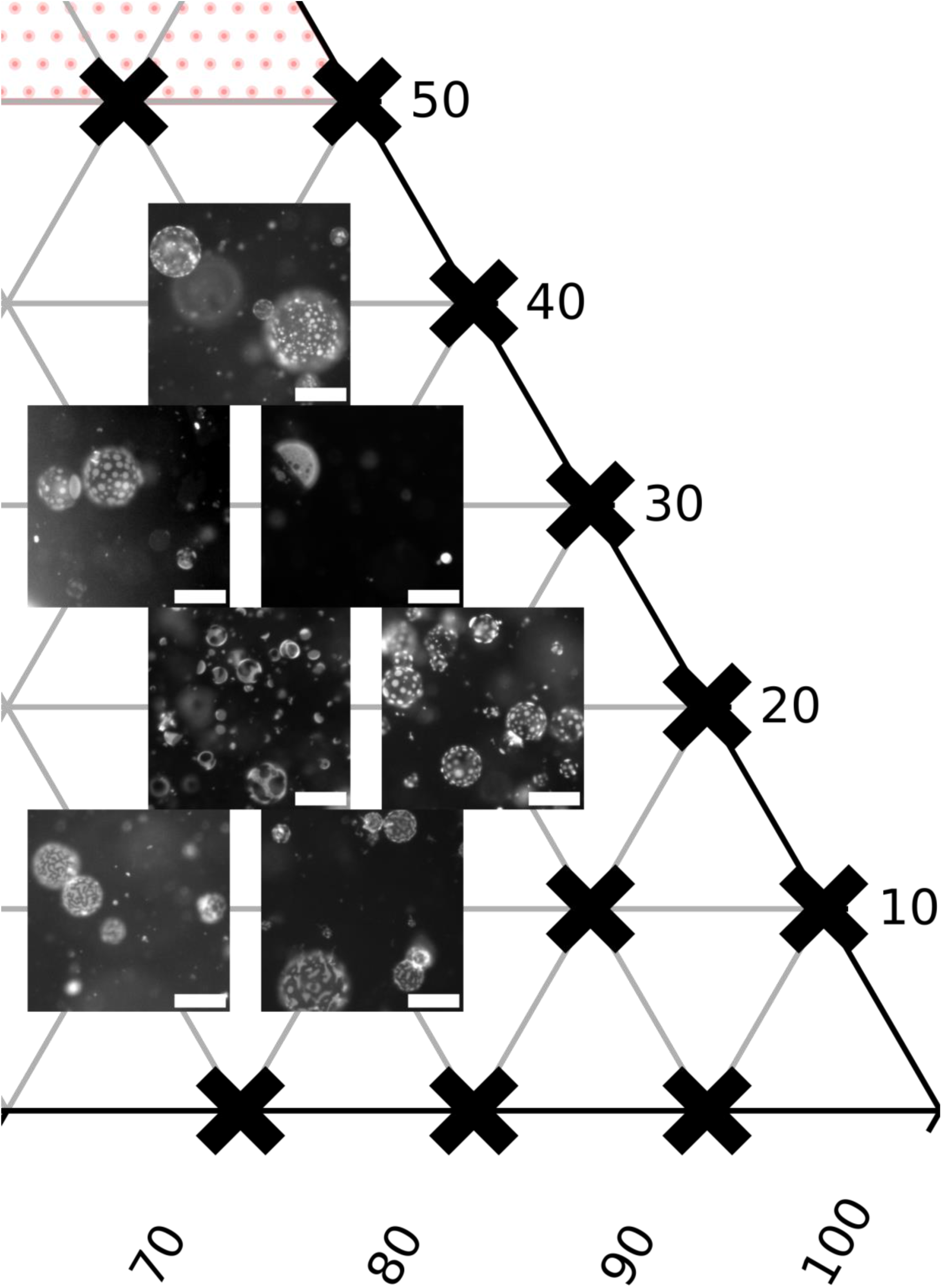
Representative fields of view for vesicles of DiphyPC/DPPE/cholesterol, covering the right third of the phase diagram in Figure 4b of the main text. The vertex at the right represents 100% DPPE, and high cholesterol is toward the top. Scalebars are 50 µm.

**Figure S12.**
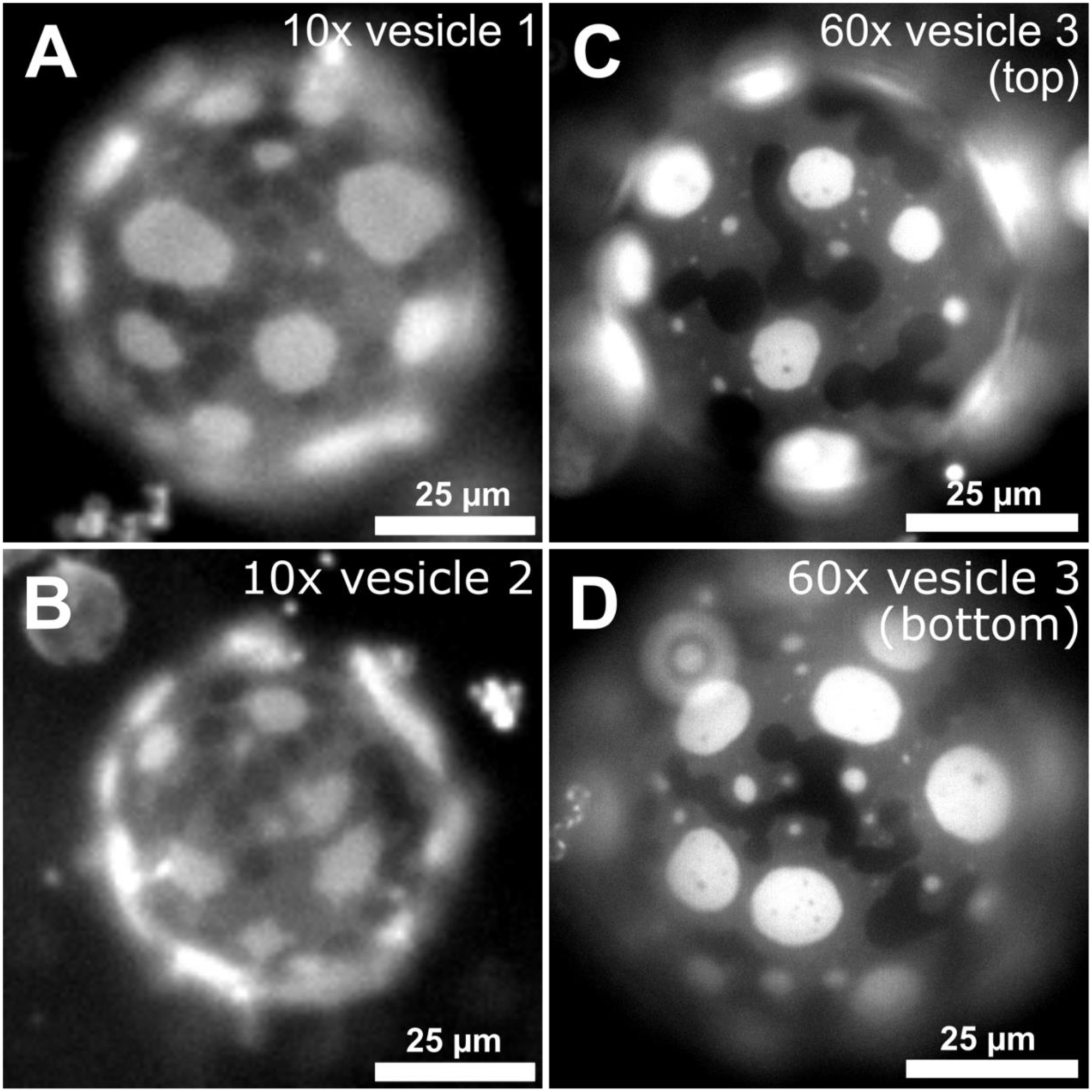
Images of vesicles composed of 10:70:20 DiphyPC/DPPE/cholesterol with three fluorescence intensities due to 3-phase coexistence of L_d_-phase domains (bright), a L_o_-phase domains (middle gray), and solid-phase domains (dark). In all cases, vesicles are close to a transition temperature in which vesicles with S/L coexistence were cooled into a region of S/L_d_/L_o_ coexistence. Shapes of the L_d_ domains are not static; they are typically noncircular due to low line tension, but they can also be noncircular due to constraints imposed by neighboring solid domains. A-B) Images of two different vesicles, collected with a 10x objective. C-D) Images of the top and bottom hemispheres of a single vesicle exhibiting three distinct fluorescence levels for coexisting L_o_, L_d_, and solid phases. As would be expected, images collected with a 60x objective are at higher resolution than those collected with a 10x objective, and it is easier to distinguish the three phases.

**Figure S13.**
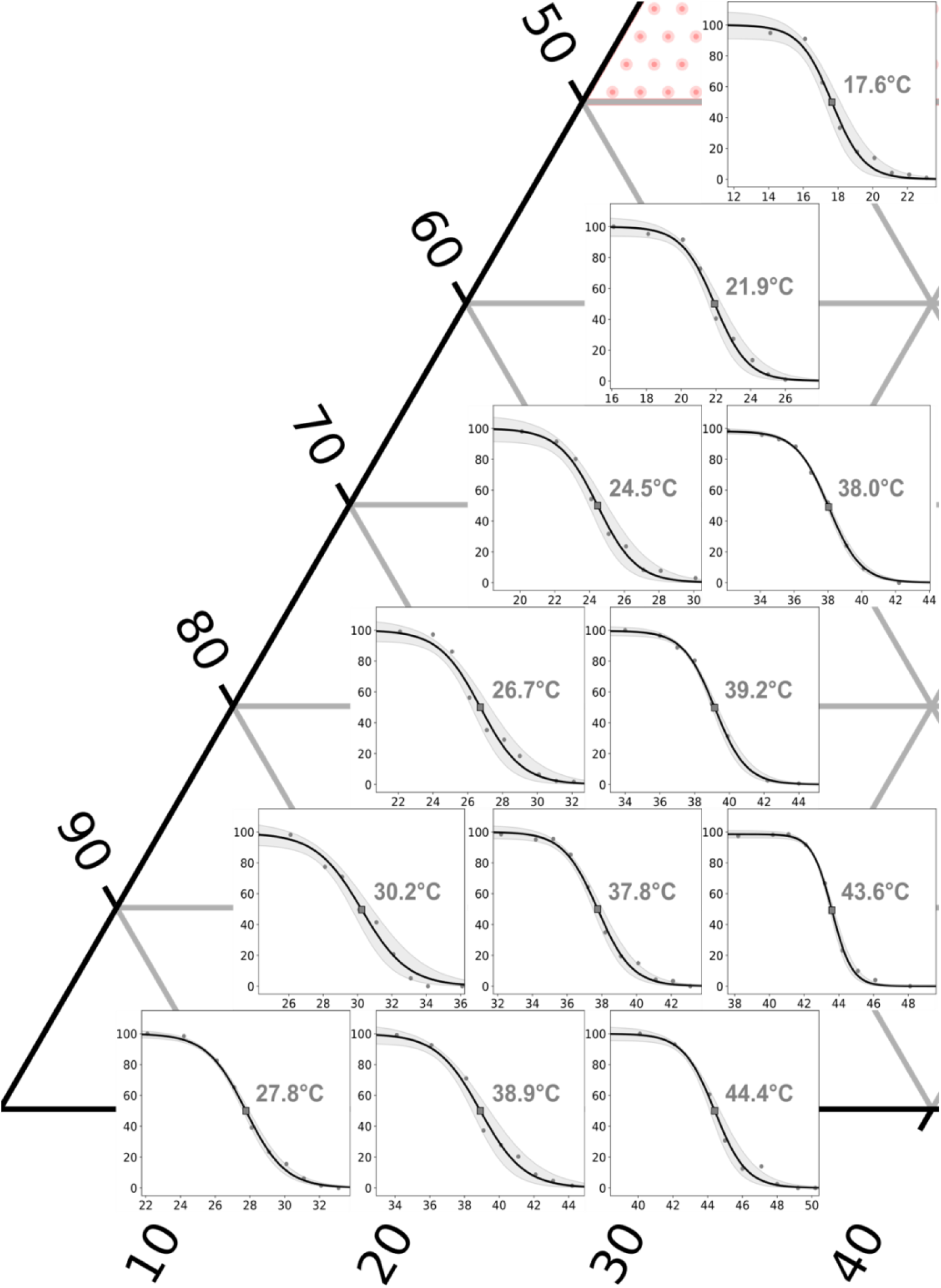
Sigmoidal fits with 95% confidence intervals (gray bands) for *T*_mix_ for vesicles of DOPC/DPPE/cholesterol, covering the left third of the phase diagram in Figure 3b of the main text. The vertex at the left represents 100% DOPC, and high cholesterol is toward the top. X-axes show temperatures (°C), y-axes show percent of phase-separated vesicles. Transition temperatures shown in gray font are transitions from one liquid phase at a high temperature to coexisting solid and liquid phases at a lower temperature.

**Figure S14.**
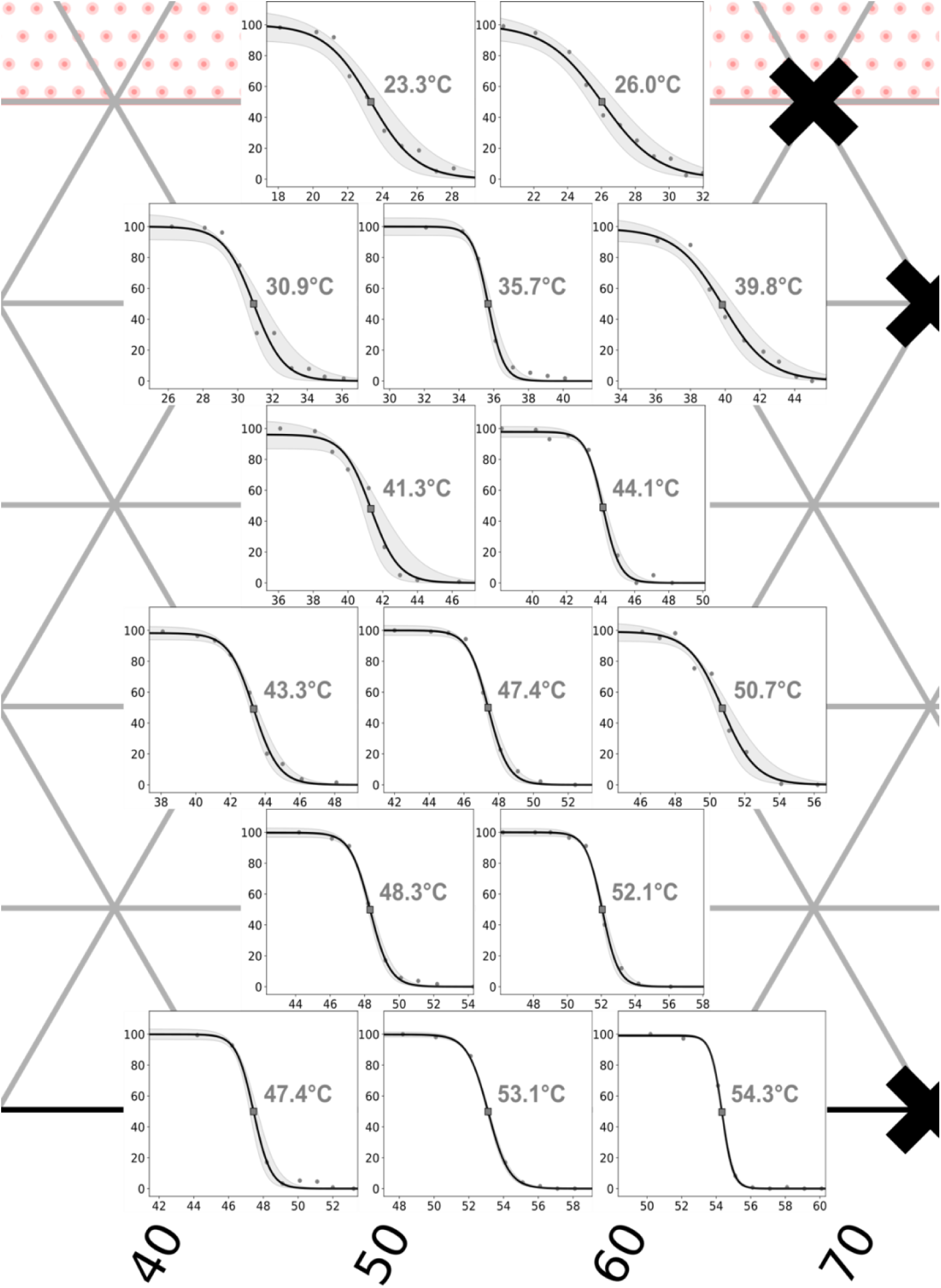
Sigmoidal fits with 95% confidence intervals (gray bands) for vesicles of DOPC/DPPE/cholesterol, covering the middle third of the phase diagram in Figure 3b of the main text. High DOPC fractions are on the left, high DPPE fractions on the right, and high cholesterol toward the top. X-axes show temperatures (°C), y-axes show percent of phase-separated vesicles. Transition temperatures shown in gray font are transitions from one liquid phase at a high temperature to coexisting solid and liquid phases at a lower temperature.

**Figure S15.**
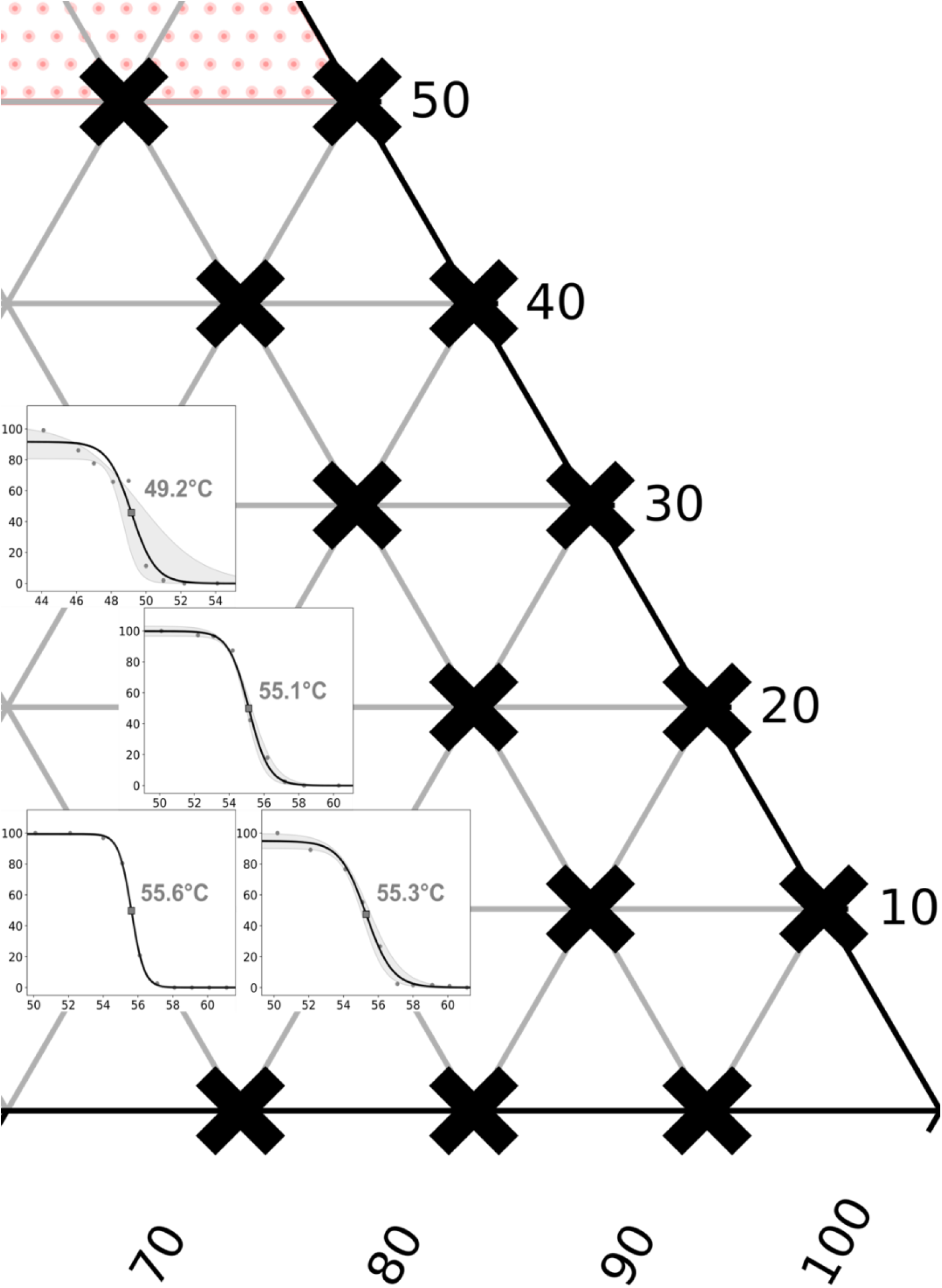
Sigmoidal fits with 95% confidence intervals (gray bands) for vesicles of DOPC/DPPE/cholesterol, covering the right third of the phase diagram in Figure 3b of the main text. The vertex at the right represents 100% DPPE, and high cholesterol is toward the top. X-axes show temperatures (°C), y-axes show percent of phase-separated vesicles. Transition temperatures shown in gray font are transitions from one liquid phase at a high temperature to coexisting solid and liquid phases at a lower temperature.

**Figure S16.**
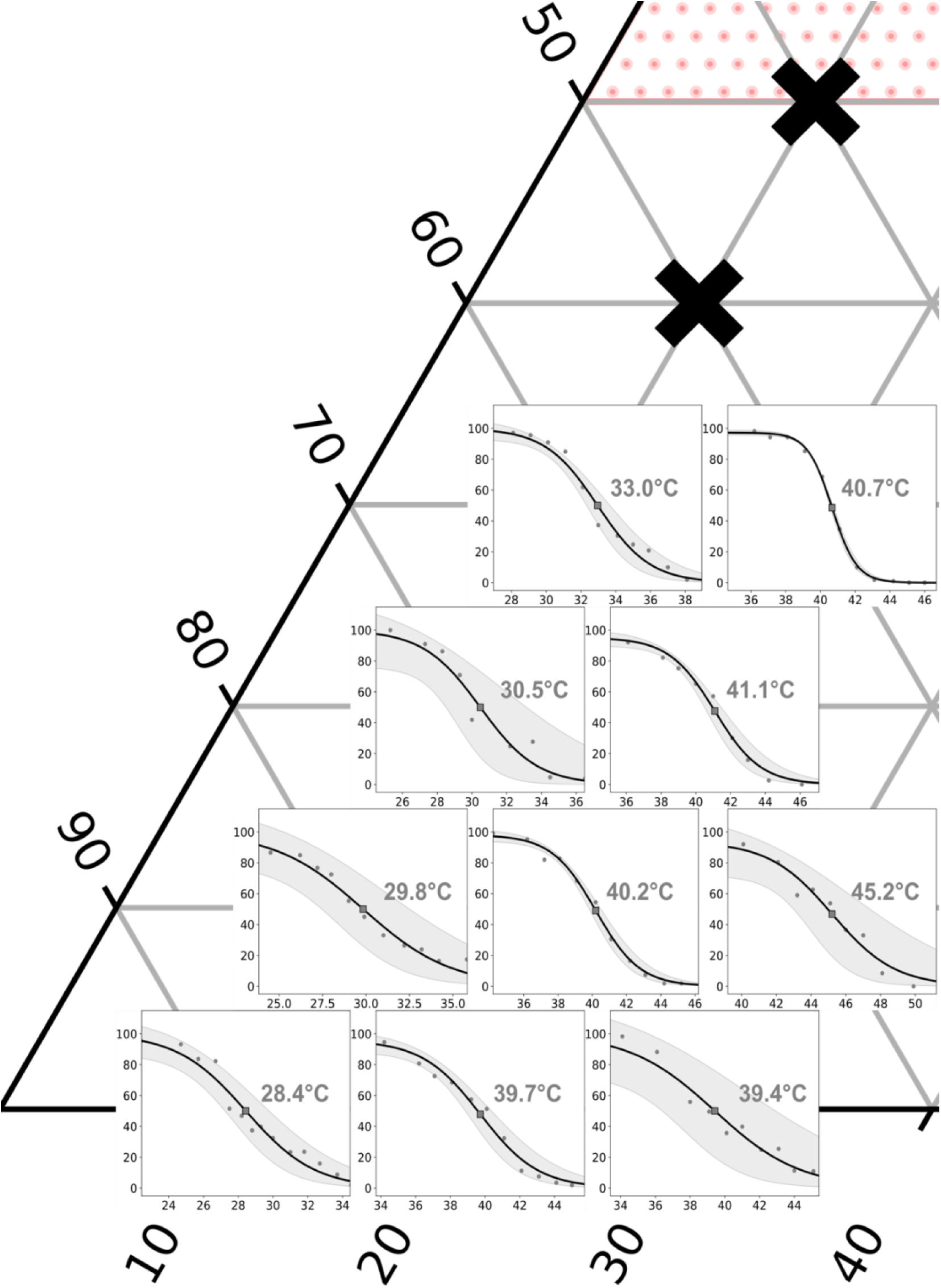
Sigmoidal fits with 95% confidence intervals (gray bands) for vesicles of DiphyPC/DPPE/cholesterol, covering the left third of the phase diagram in Figure 4b of the main text. The vertex at the left represents 100% DiphyPC, and high cholesterol is toward the top. X-axes show temperatures (°C), y-axes show percent of phase-separated vesicles. Transition temperatures shown in gray font are transitions from one liquid phase at a high temperature to coexisting solid and liquid phases at a lower temperature.

**Figure S17.**
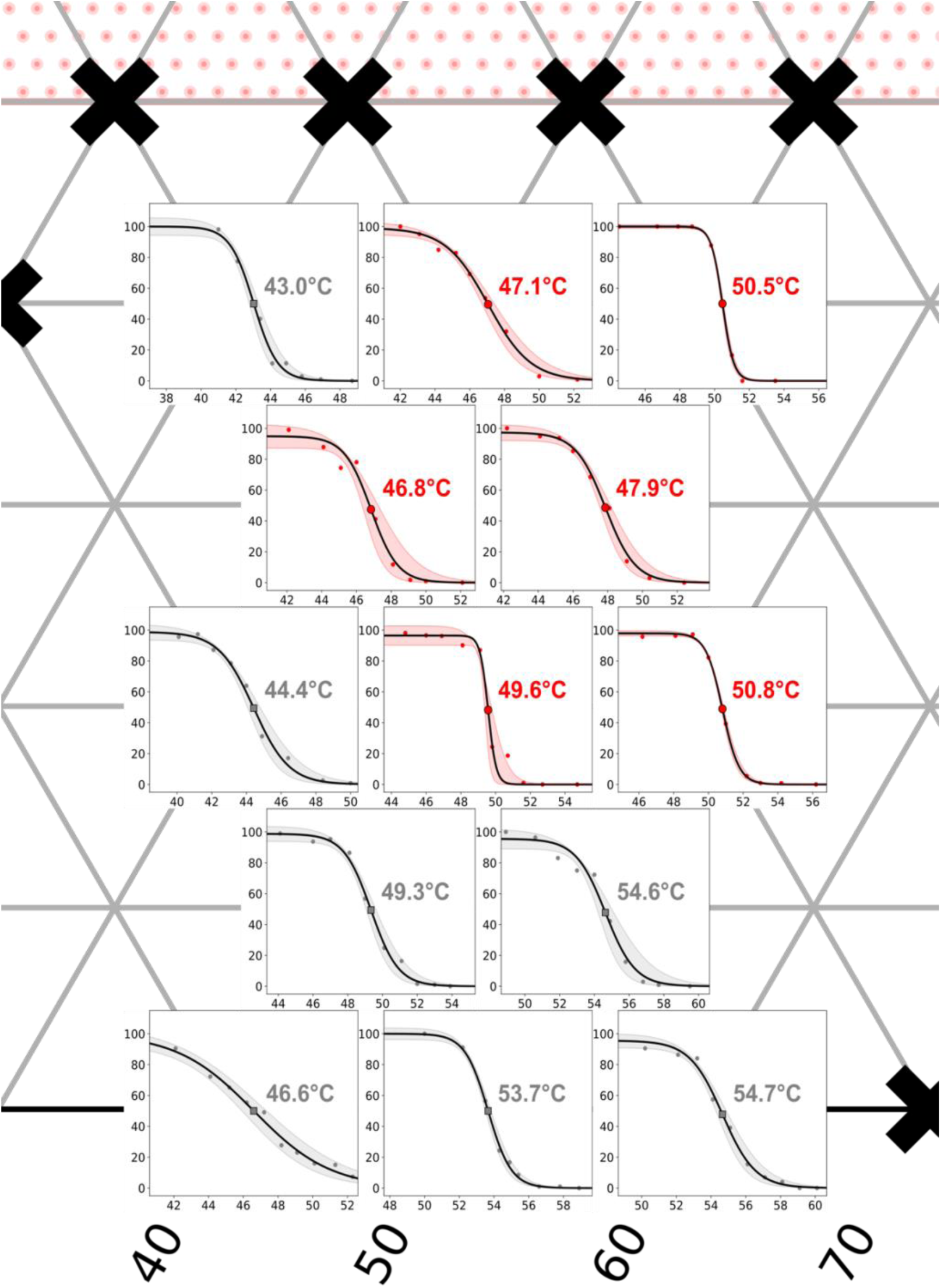
Sigmoidal fits with 95% confidence intervals (red and gray bands) for vesicles of DiphyPC/DPPE/cholesterol, covering the middle third of the phase diagram in Figure 4b of the main text. High DiphyPC fractions are on the left, high DPPE fractions on the right, and high cholesterol toward the top. X-axes show temperatures (°C), y-axes show percent of phase-separated vesicles. Transition temperatures shown in gray font are transitions from one liquid phase at a high temperature to coexisting solid and liquid phases at a lower temperature. Transition temperatures shown in red font are transitions in which coexisting L_o_ and L_d_ phases appear as temperature decreases.

**Figure S18.**
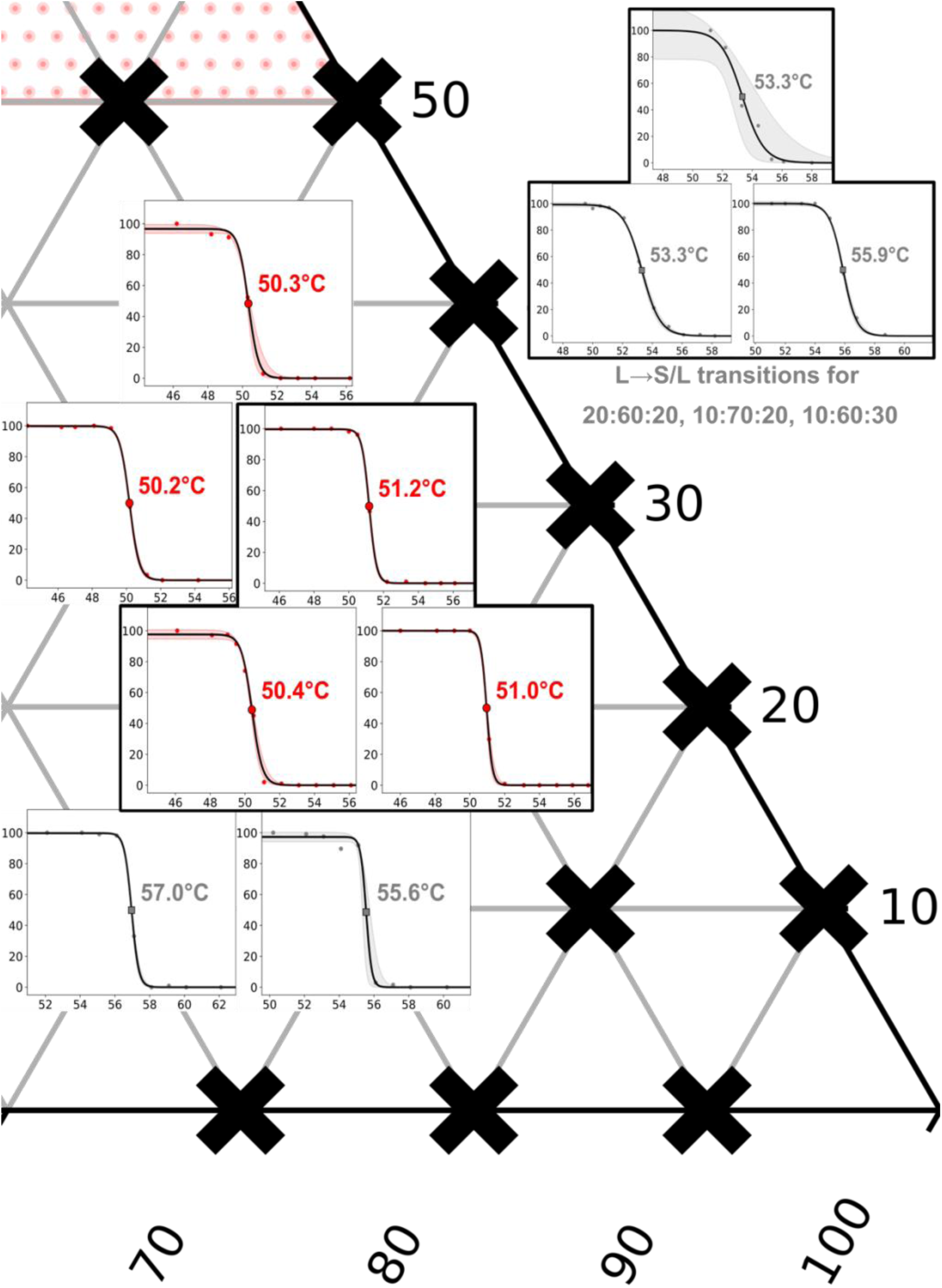
Sigmoidal fits with 95% confidence intervals (red and gray bands) for vesicles of DiphyPC/DPPE/cholesterol, covering the right third of the phase diagram as in Figure 4b of the main text. The vertex at the right represents 100% DPPE, and high cholesterol is toward the top. X-axes show temperatures (°C), y-axes show percent of phase-separated vesicles. Transition temperatures shown in gray font are transitions from one liquid phase at a high temperature to coexisting solid and liquid phases at a lower temperature. Transition temperatures shown in red font are transitions in which coexisting L_o_ and L_d_ phases appear as temperature decreases.

**Figure S19.**
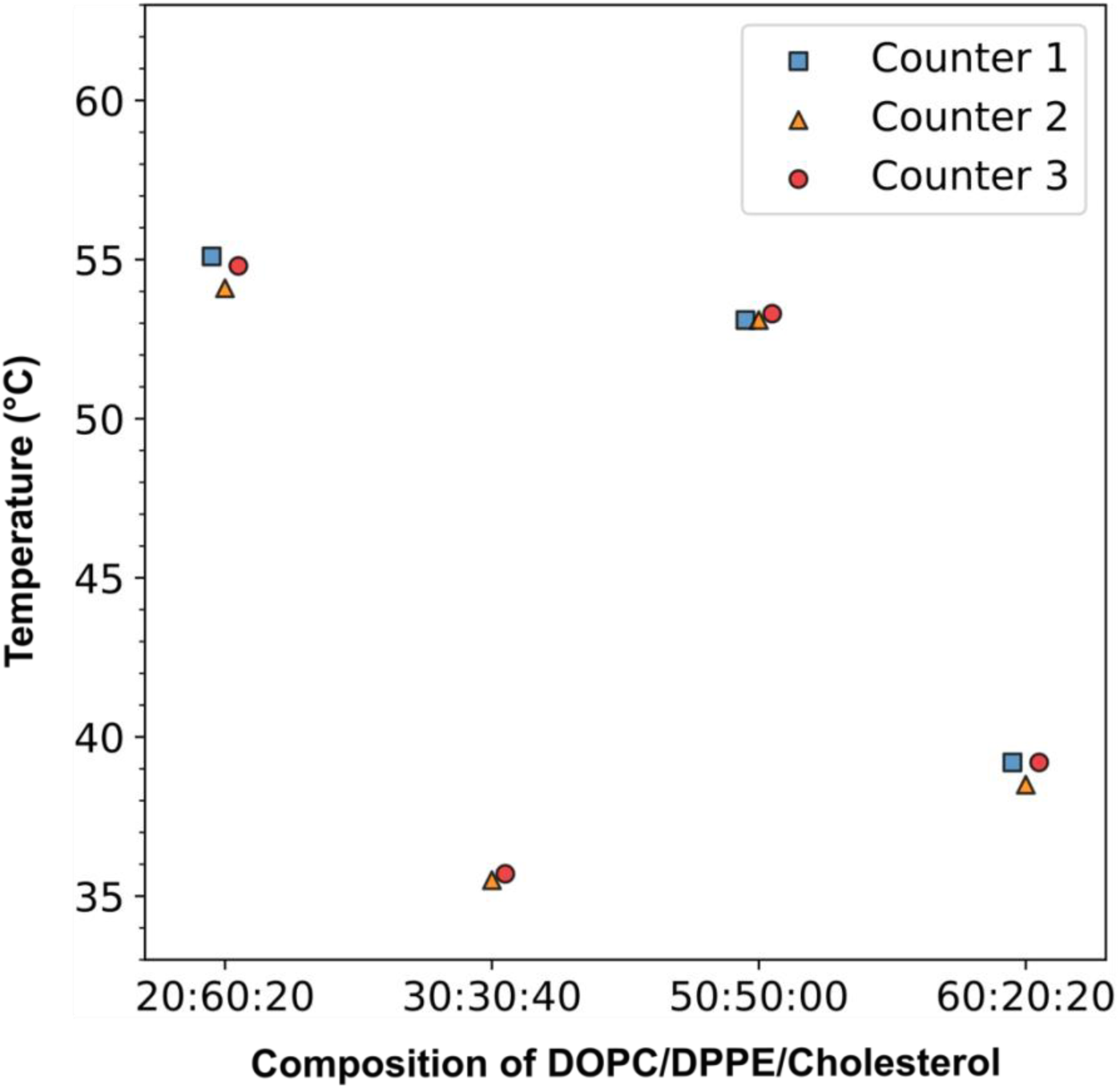
Values of *T*_mix_ are independent of which researcher analyzed the data. In this manuscript, three different researchers tabulated numbers of phase-separated vesicles from fields of view as in Fig. S4-6. To estimate the error that this introduced, all three researchers (represented by the square, triangle, and circle for Counters 1, 2, and 3) tabulated data from the same fields of view for vesicles made from the four compositions shown on the x-axis. In each case, the analyzed values of *T*_mix_ typically differed by ±0.5°C or less. For comparison, sample-to-sample variations are on the order of ±0.1 to ±1.3°C (range 0.1°C to 2.8°C).

**Table S1.**
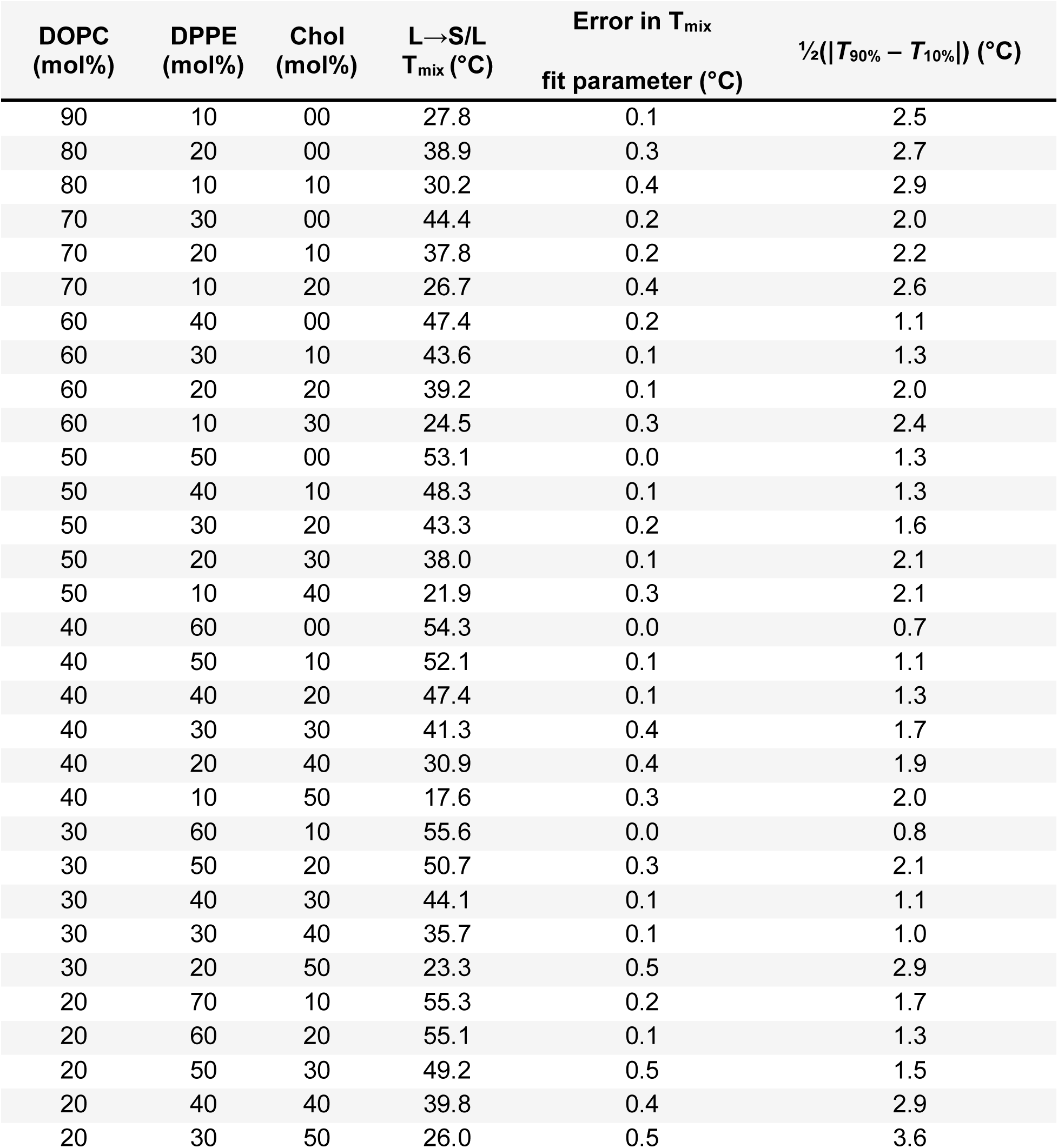
Data for transition temperatures (*T*_mix_) from one liquid phase at high temperature to coexisting solid and liquid phases at low temperature (L→S/L) for vesicles composed of varying ratios of DOPC/DPPE/cholesterol.

**Table S2.** Data for transition temperatures (*T*_mix_) from one liquid phase at high temperature to coexisting solid and liquid phases at low temperature (L→S/L) for vesicles composed of varying ratios of DiphyPC/DPPE/cholesterol. Rows in bold font indicate repeated samples from duplicate or triplicate independent experiments.

| DiphyPC<br>(mol%) | DPPE<br>(mol%) | Chol<br>(mol%) | L→S/L<br>$T_{\text{mix}}$ (°C) | Error in $T_{\text{mix}}$<br>fit parameter (°C) | $\frac{1}{2}(\vert T_{90\%} - T_{10\%}\vert )$ (°C) |
| --- | --- | --- | --- | --- | --- |
| 90 | 10 | 00 | 28.4 | 0.7 | 2.5 |
| **80** | **20** | **00** | **39.7** | **0.5** | **2.7** |
| **80** | **20** | **00** | **38.6** | **0.1** | **2.9** |
| 80 | 10 | 10 | 29.8 | 1.4 | 2.0 |
| **70** | **30** | **00** | **43.4** | **0.0** | **2.2** |
| **70** | **30** | **00** | **39.4** | **1.7** | **2.6** |
| 70 | 20 | 10 | 40.2 | 0.2 | 1.1 |
| 70 | 10 | 20 | 30.5 | 1.3 | 1.3 |
| 60 | 40 | 00 | 46.6 | 0.4 | 2.0 |
| 60 | 30 | 10 | 45.2 | 1.2 | 2.4 |
| 60 | 20 | 20 | 41.1 | 0.3 | 1.3 |
| 60 | 10 | 30 | 33.0 | 0.4 | 1.3 |
| 50 | 50 | 00 | 53.7 | 0.1 | 1.6 |
| 50 | 40 | 10 | 49.3 | 0.2 | 2.1 |
| **50** | **30** | **20** | **44.5** | **0.3** | **2.1** |
| **50** | **30** | **20** | **44.4** | **0.2** | **0.7** |
| 50 | 20 | 30 | 40.7 | 0.1 | 1.1 |
| 40 | 60 | 00 | 54.7 | 0.2 | 1.3 |
| 40 | 50 | 10 | 54.6 | 0.3 | 1.7 |
| 40 | 20 | 40 | 43.0 | 0.2 | 1.9 |
| 30 | 60 | 10 | 55.6 | 0.2 | 2.0 |
| 20 | 70 | 10 | 57.0 | 0.0 | 0.8 |
| **20** | **60** | **20** | **54.5** | **0.1** | **0.9** |
| **20** | **60** | **20** | **53.3** | **0.1** | **1.3** |
| **20** | **60** | **20** | **54.2** | **0.3** | **1.8** |
| **10** | **70** | **20** | **55.6** | **0.0** | **0.7** |
| **10** | **70** | **20** | **55.9** | **0.1** | **1.0** |
| **10** | **70** | **20** | **56.0** | **0.2** | **0.8** |
| **10** | **60** | **30** | **52.4** | **0.4** | **1.9** |
| **10** | **60** | **30** | **52.2** | **1.2** | **1.5** |
| **10** | **60** | **30** | **53.3** | **0.6** | **1.6** |

**Table S3.** Data for transition temperatures (*T*_mix_) for vesicles with a liquid phase (and perhaps a solid phase) at high temperatures, and coexisting L_o_ and L_d_ phases at lower temperatures (L→ L_o_/L_d_), for vesicles composed of varying ratios of DiphyPC/DPPE/cholesterol. Rows in bold font indicate repeated samples from duplicate or triplicate independent experiments.

| DiphyPC<br>(mol%) | DPPE<br>(mol%) | Chol<br>(mol%) | L→ L<sub>o</sub>/L<sub>d</sub><br>T<sub>mix</sub> (°C) | Error in T<sub>mix</sub><br>fit parameter (°C) | $\frac{1}{2}(\vert T_{90\%} - T_{10\%}\vert )$ (°C) |
| --- | --- | --- | --- | --- | --- |
| 40 | 40 | 20 | 49.6 | 0.2 | 2.5 |
| 40 | 30 | 30 | 46.8 | 0.3 | 2.7 |
| 30 | 50 | 20 | 50.8 | 0.1 | 2.9 |
| 30 | 40 | 30 | 47.9 | 0.2 | 2.0 |
| 30 | 30 | 40 | 47.1 | 0.2 | 2.2 |
| **20** | **60** | **20** | **49.9** | **0.1** | **2.6** |
| **20** | **60** | **20** | **50.4** | **0.1** | **1.1** |
| **20** | **60** | **20** | **51.5** | **0.2** | **1.3** |
| 20 | 50 | 30 | 50.2 | 0.0 | 2.0 |
| 20 | 40 | 40 | 50.5 | 0.1 | 2.4 |
| **10** | **70** | **20** | **51.3** | **0.0** | **1.3** |
| **10** | **70** | **20** | **51.0** | **0.0** | **1.3** |
| **10** | **70** | **20** | **51.4** | **0.1** | **1.6** |
| **10** | **60** | **30** | **51.1** | **0.2** | **2.1** |
| **10** | **60** | **30** | **51.2** | **0.0** | **2.1** |
| **10** | **60** | **30** | **51.3** | **0.0** | **0.7** |
| 10 | 50 | 40 | 50.3 | 0.1 | 1.1 |

**Table S4.**
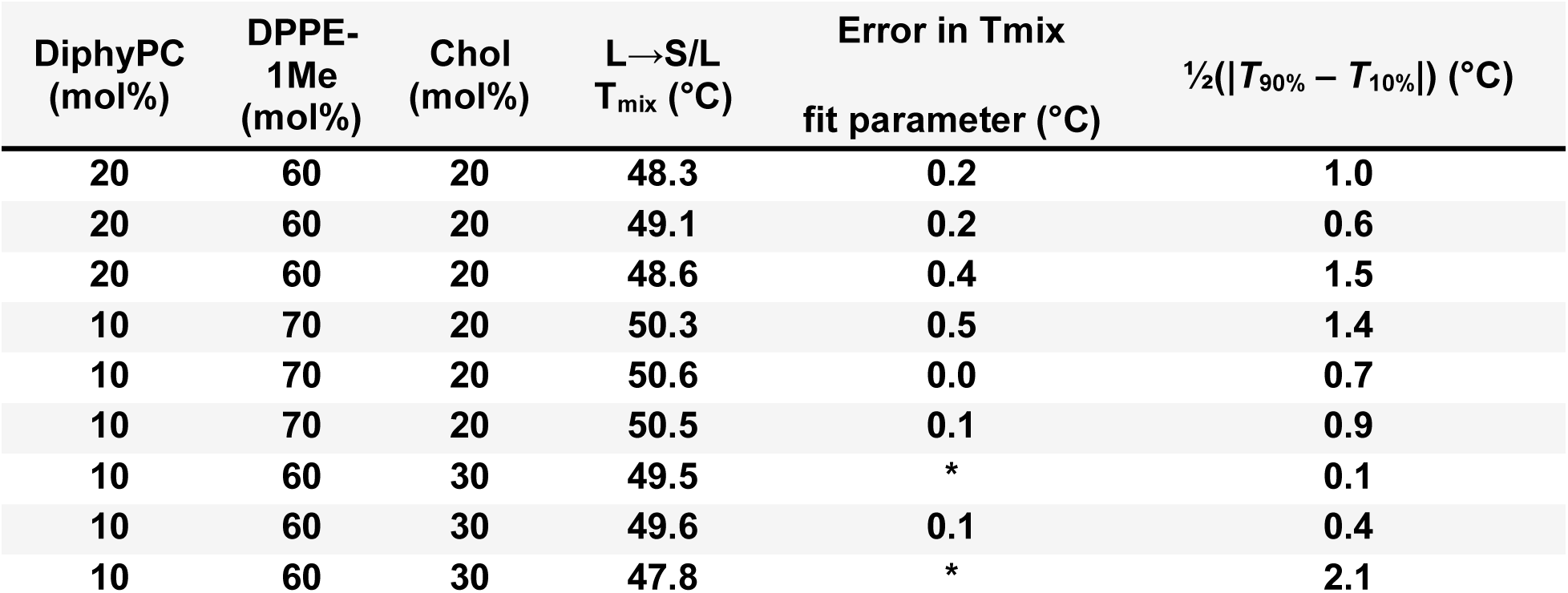
Data for transition temperatures (*T*_mix_) from one liquid phase at high temperature to coexisting solid and liquid phases at low temperature (L→S/L) for vesicles composed of varying ratios of DiphyPC/DPPE-1Me/cholesterol. Rows in bold font indicate repeated samples from duplicate or triplicate independent experiments. Star symbols (**\***) indicate *T*_mix_ fit parameter values that were non-physical due to the proximity of the transition to the 3-phase region at temperatures just below the L→S/L *T*_mix_. The temperature range to observe S/L coexistence narrows with increasing headgroup methylation and increasing DOPC composition.

| DiphyPC<br>(mol%) | DPPE-1Me<br>(mol%) | Chol<br>(mol%) | L→S/L<br>$T_{\text{mix}}$ (°C) | Error in $T_{\text{mix}}$<br>fit parameter (°C) | $\frac{1}{2}(\vert T_{90\%} - T_{10\%}\vert )$ (°C) |
| --- | --- | --- | --- | --- | --- |
| **20** | **60** | **20** | **48.3** | **0.2** | **1.0** |
| **20** | **60** | **20** | **49.1** | **0.2** | **0.6** |
| **20** | **60** | **20** | **48.6** | **0.4** | **1.5** |
| **10** | **70** | **20** | **50.3** | **0.5** | **1.4** |
| **10** | **70** | **20** | **50.6** | **0.0** | **0.7** |
| **10** | **70** | **20** | **50.5** | **0.1** | **0.9** |
| **10** | **60** | **30** | **49.5** | * | **0.1** |
| **10** | **60** | **30** | **49.6** | **0.1** | **0.4** |
| **10** | **60** | **30** | **47.8** | * | **2.1** |

**Table S5.** Data for transition temperatures (*T*_mix_) for vesicles with a liquid phase (and perhaps a solid phase) at high temperatures, and coexisting L_o_ and L_d_ phases at lower temperatures (L→ L_o_/L_d_), for vesicles composed of varying ratios of DiphyPC/DPPE-1Me/cholesterol. Rows in bold font indicate repeated samples from duplicate or triplicate independent experiments.

| DiphyPC<br>(mol%) | DPPE-1Me<br>(mol%) | Chol<br>(mol%) | L→ L<sub>o</sub>/L<sub>d</sub><br>$T_{\text{mix}}$ (°C) | Error in $T_{\text{mix}}$<br>fit parameter (°C) | $\frac{1}{2}(\vert T_{90\%} - T_{10\%}\vert )$ (°C) |
| --- | --- | --- | --- | --- | --- |
| **20** | **60** | **20** | **48.8** | **0.1** | **0.4** |
| **20** | **60** | **20** | **48.8** | **0.1** | **0.5** |
| **20** | **60** | **20** | **48.6** | **0.0** | **0.4** |
| **10** | **70** | **20** | **49.4** | **0.0** | **0.2** |
| **10** | **70** | **20** | **49.2** | **0.0** | **0.4** |
| **10** | **70** | **20** | **49.9** | **0.1** | **0.6** |
| **10** | **60** | **30** | **49.5** | **0.0** | **0.3** |
| **10** | **60** | **30** | **49.3** | **0.0** | **0.3** |
| **10** | **60** | **30** | **49.7** | **0.0** | **0.3** |

**Table S6.** Data for transition temperatures (*T*_mix_) from one liquid phase at high temperature to coexisting solid and liquid phases at low temperature (L→S/L) for vesicles composed of varying ratios of DiphyPC/DPPE-2Me/cholesterol. Rows in bold font indicate repeated samples from duplicate or triplicate independent experiments.

| DiphyPC<br>(mol%) | DPPE-2Me<br>(mol%) | Chol<br>(mol%) | $T_{\text{mix}}$ (°C) | Error in $T_{\text{mix}}$<br>fit parameter (°C) | $\frac{1}{2}(\vert T_{90\%} - T_{10\%}\vert )$ (°C) |
| --- | --- | --- | --- | --- | --- |
| **10** | **70** | **20** | **46.7** | **0.1** | **0.5** |
| **10** | **70** | **20** | **44.7** | **0.1** | **0.5** |
| **10** | **70** | **20** | **45.5** | **0.1** | **0.6** |

**Table S7.** Data for transition temperatures (*T*_mix_) for vesicles with a liquid phase (and perhaps a solid phase) at high temperatures, and coexisting L_o_ and L_d_ phases at lower temperatures (L→ L_o_/L_d_), for vesicles composed of varying ratios of DiphyPC/DPPE-2Me/cholesterol. Rows in bold font indicate repeated samples from duplicate or triplicate independent experiments.

| DiphyPC<br>(mol%) | DPPE-2Me<br>(mol%) | Chol<br>(mol%) | $L \rightarrow L_o/L_d$<br>$T_{\text{mix}}$ (°C) | Error in $T_{\text{mix}}$<br>fit parameter (°C) | $\frac{1}{2}(\vert T_{90\%} - T_{10\%}\vert )$ (°C) |
| --- | --- | --- | --- | --- | --- |
| **20** | **60** | **20** | **45.8** | **0.0** | **0.6** |
| **20** | **60** | **20** | **45.2** | **0.1** | **0.8** |
| **20** | **60** | **20** | **46.3** | **0.0** | **0.4** |
| **10** | **70** | **20** | **44.4** | **0.1** | **0.7** |
| **10** | **70** | **20** | **46.3** | **0.1** | **0.3** |
| **10** | **70** | **20** | **45.3** | **0.0** | **0.6** |
| **10** | **60** | **30** | **47.2** | **0.0** | **1.1** |
| **10** | **60** | **30** | **47.3** | **0.1** | **0.3** |
| **10** | **60** | **30** | **46.3** | **0.1** | **0.9** |

**Table S8.** Data for transition temperatures (*T*_mix_) for vesicles with a liquid phase (and perhaps a solid phase) at high temperatures, and coexisting L_o_ and L_d_ phases at lower temperatures (L→ L_o_/L_d_), for vesicles composed of varying ratios of DiphyPC/DPPC/cholesterol. Rows in bold font indicate repeated samples from duplicate or triplicate independent experiments.

| DiphyPC<br>(mol%) | DPPC<br>(mol%) | Chol<br>(mol%) | $L \rightarrow L_o/L_d$<br>$T_{mix}$ (°C) | Error in $T_{mix}$<br>fit parameter (°C) | $\frac{1}{2}(\vert T_{90\%} - T_{10\%}\vert )$ (°C) |
| --- | --- | --- | --- | --- | --- |
| **20** | **60** | **20** | **41.4** | **0.4** | **2.0** |
| **20** | **60** | **20** | **42.9** | **0.3** | **2.3** |
| **20** | **60** | **20** | **40.8** | **0.3** | **2.4** |
| **10** | **70** | **20** | **40.4** | **0.2** | **1.1** |
| **10** | **70** | **20** | **38.5** | **0.4** | **2.5** |
| **10** | **70** | **20** | **39.1** | **0.2** | **2.1** |
| **10** | **60** | **30** | **43.0** | **0.2** | **2.2** |
| **10** | **60** | **30** | **43.8** | **0.3** | **1.9** |
| **10** | **60** | **30** | **44.8** | **0.1** | **1.8** |

## REFERENCES

1. Leveille, C.L., C.E. Cornell, A.J. Merz, and S.L. Keller. 2022. Yeast cells actively tune their membranes to phase separate at temperatures that scale with growth temperatures. Proc. Natl. Acad. Sci. USA. 119:e2116007119.

2. Burns, M., K. Wisser, J. Wu, I. Levental, and S.L. Veatch. 2017. Miscibility Transition Temperature Scales with Growth Temperature in a Zebrafish Cell Line. Biophys. J. 113:1212–1222.

3. Lorent, J.H., K.R. Levental, L. Ganesan, G. Rivera-Longsworth, E. Sezgin, M. Doktorova, E. Lyman, and I. Levental. 2020. Plasma membranes are asymmetric in lipid unsaturation, packing and protein shape. Nat. Chem. Biol. 16:644–652.

4. Reinhard, J., C.L. Leveille, C.E. Cornell, A.J. Merz, C. Klose, R. Ernst, and S.L. Keller. 2023. Remodeling of yeast vacuole membrane lipidomes from the log (one phase) to stationary stage (two phases). Biophys. J. 122:1043–1057.

5. Kim, H., and I. Budin. 2024. Intracellular sphingolipid sorting drives membrane phase separation in the yeast vacuole. J. Biol. Chem. 300:105496.

6. Wilson, K.J., H.Q. Nguyen, J. Gervay-Hague, and S.L. Keller. 2024. Sterol–lipids enable large-scale, liquid–liquid phase separation in bilayer membranes of only two components. Proc. Natl. Acad. Sci. USA. 121:e2401241121.

7. Dietrich, C., L.A. Bagatolli, Z.N. Volovyk, N.L. Thompson, M. Levi, K. Jacobson, and E. Gratton. 2001. Lipid Rafts Reconstituted in Model Membranes. Biophys. J. 80:1417–1428.

8. Samsonov, A.V., I. Mihalyov, and F.S. Cohen. 2001. Characterization of Cholesterol-Sphingomyelin Domains and Their Dynamics in Bilayer Membranes. Biophys. J. 81:1486– 1500.

9. Veatch, S.L., and S.L. Keller. 2002. Organization in Lipid Membranes Containing Cholesterol. Phys. Rev. Lett. 89:268101.

10. Koynova, R., and M. Caffrey. 1998. Phases and phase transitions of the phosphatidylcholines. BBA - Rev. Biomembr. 1376:91–145.

11. Veatch, S.L., and S.L. Keller. 2005. Miscibility Phase Diagrams of Giant Vesicles Containing Sphingomyelin. Phys. Rev. Lett. 94:148101.

12. Ionova, I.V., V.A. Livshits, and D. Marsh. 2012. Phase Diagram of Ternary Cholesterol/Palmitoylsphingomyelin/Palmitoyloleoyl-Phosphatidylcholine Mixtures: Spin-Label EPR Study of Lipid-Raft Formation. Biophys. J. 102:1856–1865.

13. Lin, W.-C., C.D. Blanchette, and M.L. Longo. 2007. Fluid-Phase Chain Unsaturation Controlling Domain Microstructure and Phase in Ternary Lipid Bilayers Containing GalCer and Cholesterol. Biophys. J. 92:2831–2841.

14. Castro, B.M., L.C. Silva, A. Fedorov, R.F.M. de Almeida, and M. Prieto. 2009. Cholesterol-rich Fluid Membranes Solubilize Ceramide Domains: Implications for the Structure and Dynamics of Mammalian Intracellular and Plasma Membranes. J. Biol. Chem. 284:22978– 22987.

15. Blosser, M.C., J.B. Starr, C.W. Turtle, J. Ashcraft, and S.L. Keller. 2013. Minimal Effect of Lipid Charge on Membrane Miscibility Phase Behavior in Three Ternary Systems. Biophys. J. 104:2629–2638.

16. Vequi-Suplicy, C.C., K.A. Riske, R.L. Knorr, and R. Dimova. 2010. Vesicles with charged domains. BBA - Biomembr. 1798:1338–1347.

17. Pitman, M.C., F. Suits, K. Gawrisch, and S.E. Feller. 2005. Molecular dynamics investigation of dynamical properties of phosphatidylethanolamine lipid bilayers. J. Chem. Phys. 122:244715.

18. Suits, F., M.C. Pitman, and S.E. Feller. 2005. Molecular dynamics investigation of the structural properties of phosphatidylethanolamine lipid bilayers. J. Chem. Phys. 122:244714.

19. Cullis, P.R., M.J. Hope, and C.P.S. Tilcock. 1986. Lipid polymorphism and the roles of lipids in membranes. Chem. Phys. Lipids. 40:127–144.

20. Davies, M.A., W. Hubner, A. Blume, and R. Mendelsohn. 1992. Acyl chain conformational ordering in 1,2 dipalmitoylphosphatidylethanolamine. Integration of FT-IR and 2H NMR results. Biophys. J. 63:1059–1062.

21. Thurmond, R.L., S.W. Dodd, and M.F. Brown. 1991. Molecular areas of phospholipids as determined by 2H NMR spectroscopy. Comparison of phosphatidylethanolamines and phosphatidylcholines. Biophys. J. 59:108–113.

22. Koynova, R., and M. Caffrey. 1994. Phases and phase transitions of the hydrated phosphatidylethanolamines. Chem. Phys. Lipids. 69:1–34.

23. Keller, S.L., S.M. Bezrukov, S.M. Gruner, M.W. Tate, I. Vodyanoy, and V.A. Parsegian. 1993. Probability of alamethicin conductance states varies with nonlamellar tendency of bilayer phospholipids. Biophys. J. 65:23–27.

24. Brown, M.F. 1994. Modulation of rhodopsin function by properties of the membrane bilayer. Chem. Phys. Lipids. 73:159–180.

25. Bogdanov, M., and W. Dowhan. 1998. Phospholipid-assisted protein folding: phosphatidylethanolamine is required at a late step of the conformational maturation of the polytopic membrane protein lactose permease. EMBO J. 17:5255–5264.

26. Wikström, M., A.A. Kelly, A. Georgiev, H.M. Eriksson, M.R. Klement, M. Bogdanov, W. Dowhan, and Å. Wieslander. 2009. Lipid-engineered *Escherichia coli* Membranes Reveal Critical Lipid Headgroup Size for Protein Function. J. Biol. Chem. 284:954–965.

27. Andersen, O.S., and R.E. Koeppe 2nd. 2007. Bilayer Thickness and Membrane Protein Function: An Energetic Perspective. *An*nu. Rev. Biophys. 36:107–130.

28. Siegel, D.P. 1993. Energetics of intermediates in membrane fusion: comparison of stalk and inverted micellar intermediate mechanisms. Biophys. J. 65:2124–2140.

29. Allender, D.W., and M. Schick. 2023. On the force between “rafts.” Eur. Phys. J. E. 46:85.

30. Cornell, C.E., A.D. Skinkle, S. He, I. Levental, K.R. Levental, and S.L. Keller. 2018. Tuning Length Scales of Small Domains in Cell-Derived Membranes and Synthetic Model Membranes. Biophys. J. 115:690–701.

31. Shyamsunder, E., S.M. Gruner, M.W. Tate, D.C. Turner, P.T.C. So, and C.P.S. Tilcock. 1988. Observation of inverted cubic phase in hydrated dioleoylphosphatidylethanolamine membranes. Biochem. 27:2332–2336.

32. So, P.T.C., S.M. Gruner, and S. Erramilli. 1993. Pressure-induced topological phase transitions in membranes. Phys. Rev. Lett. 70:3455–3458.

33. Winnikoff, J.R., D. Milshteyn, S.J. Vargas-Urbano, M.A. Pedraza-Joya, A.M. Armando, O. Quehenberger, A. Sodt, R.E. Gillilan, E.A. Dennis, E. Lyman, S.H.D. Haddock, and I. Budin. 2024. Homeocurvature adaptation of phospholipids to pressure in deep-sea invertebrates. Science. 384:1482–1488.

34. Veatch, S.L., and S.L. Keller. 2003. Separation of Liquid Phases in Giant Vesicles of Ternary Mixtures of Phospholipids and Cholesterol. Biophys. J. 85:3074–3083.

35. Veatch, S.L., K. Gawrisch, and S.L. Keller. 2006. Closed-Loop Miscibility Gap and Quantitative Tie-Lines in Ternary Membranes Containing Diphytanoyl PC. Biophys. J. 90:4428–4436.

36. Garbès Putzel, G., and M. Schick. 2008. Phenomenological Model and Phase Behavior of Saturated and Unsaturated Lipids and Cholesterol. Biophys. J. 95:4756–4762.

37. Klose, C., M.A. Surma, M.J. Gerl, F. Meyenhofer, A. Shevchenko, and K. Simons. 2012. Flexibility of a Eukaryotic Lipidome – Insights from Yeast Lipidomics. PLOS ONE. 7:e35063.

38. Marrink, S.J., V. Corradi, P.C.T. Souza, H.I. Ingólfsson, D.P. Tieleman, and M.S.P. Sansom. 2019. Computational Modeling of Realistic Cell Membranes. Chem. Rev. 119:6184–6226.

39. Hossein, A., and M. Deserno. 2020. Spontaneous Curvature, Differential Stress, and Bending Modulus of Asymmetric Lipid Membranes. Biophys. J. 118:624–642.

40. Keller, S.L., S.M. Gruner, and K. Gawrisch. 1996. Small concentrations of alamethicin induce a cubic phase in bulk phosphatidylethanolamine mixtures. BBA - Biomembr. 1278:241–246.

41. Lindsey, H., N.O. Petersen, and S.I. Chan. 1979. Physicochemical characterization of 1,2-diphytanoyl-*sn*-glycero-3-phosphocholine in model membrane systems. BBA - Biomembr. 555:147–167.

42. Veatch, S.L. 2007. Electro-formation and fluorescence microscopy of giant vesicles with coexisting liquid phases. Methods Mol. Biol. 398:59–72.

43. Veatch, S.L., and S.L. Keller. 2005. Seeing spots: Complex phase behavior in simple membranes. *BBA - Mol*. Cell Res. 1746:172–185.

44. Weakly, H.M.J., K.J. Wilson, G.J. Goetz, E.L. Pruitt, A. Li, L. Xu, and S.L. Keller. 2024. Several common methods of making vesicles (except an emulsion method) capture intended lipid ratios. Biophys. J. 123:3452–3462.

45. Schindelin, J., I. Arganda-Carreras, E. Frise, V. Kaynig, M. Longair, T. Pietzsch, S. Preibisch, C. Rueden, S. Saalfeld, B. Schmid, J.-Y. Tinevez, D.J. White, V. Hartenstein, K. Eliceiri, P. Tomancak, and A. Cardona. 2012. Fiji: an open-source platform for biological-image analysis. Nat. Methods. 9:676–682.

46. Virtanen, P., R. Gommers, T.E. Oliphant, M. Haberland, T. Reddy, D. Cournapeau, E. Burovski, P. Peterson, W. Weckesser, J. Bright, S.J. van der Walt, M. Brett, J. Wilson, K.J. Millman, N. Mayorov, A.R.J. Nelson, E. Jones, R. Kern, E. Larson, C.J. Carey, İ. Polat, Y. Feng, E.W. Moore, J. VanderPlas, D. Laxalde, J. Perktold, R. Cimrman, I. Henriksen, E.A. Quintero, C.R. Harris, A.M. Archibald, A.H. Ribeiro, F. Pedregosa, and P. van Mulbregt. 2020. SciPy 1.0: fundamental algorithms for scientific computing in Python. Nat. Methods. 17:261–272.

47. Hunter, J.D. 2007. Matplotlib: A 2D Graphics Environment. Comput. Sci. Eng. 9:90–95.

48. Ikeda, Y. 2023. yuzie007/mpltern: 1.0.2. Zenodo. 10.5281/zenodo.8289090.

49. Huang, J., J.T. Buboltz, and G.W. Feigenson. 1999. Maximum solubility of cholesterol in phosphatidylcholine and phosphatidylethanolamine bilayers. BBA - Biomembr. 1417:89– 100.

50. Shaikh, S.R., V. Cherezov, M. Caffrey, S.P. Soni, D. LoCascio, W. Stillwell, and S.R. Wassall. 2006. Molecular Organization of Cholesterol in Unsaturated Phosphatidylethanolamines: X-ray Diffraction and Solid State 2H NMR Reveal Differences with Phosphatidylcholines. J. Am. Chem. Soc. 128:5375–5383.

51. Sakuma, Y., M. Imai, M. Yanagisawa, and S. Komura. 2008. Adhesion of binary giant vesicles containing negative spontaneous curvature lipids induced by phase separation. Eur. Phys. J. E. 25:403–413.

52. Blume, A., and R.G. Griffin. 1982. Carbon-13 and deuterium nuclear magnetic resonance study of the interaction of cholesterol with phosphatidylethanolamine. Biochem. 21:6230– 6242.

53. Wu, S.H.W., and H.M. McConnell. 1975. Phase separations in phospholipid membranes. Biochem. 14:847–854.

54. Davis, J.H., J.J. Clair, and J. Juhasz. 2009. Phase Equilibria in DOPC/DPPC-d62/Cholesterol Mixtures. Biophys. J. 96:521–539.

55. Chen, D., and M.M. Santore. 2014. 1,2-Dipalmitoyl-sn-glycero-3-phosphocholine (DPPC)-Rich Domain Formation in Binary Phospholipid Vesicle Membranes: Two-Dimensional Nucleation and Growth. Langmuir. 30:9484–9493.

56. Wan, H., G. Jeon, W. Xin, G.M. Grason, and M.M. Santore. 2024. Flower-shaped 2D crystals grown in curved fluid vesicle membranes. Nat. Commun. 15:3442.

57. Tornabene, T.G., and T.A. Langworthy. 1979. Diphytanyl and Dibiphytanyl Glycerol Ether Lipids of Methanogenic Archaebacteria. Science. 203:51–53.

58. Pataraia, S., Y. Liu, R. Lipowsky, and R. Dimova. 2014. Effect of cytochrome c on the phase behavior of charged multicomponent lipid membranes. BBA - Biomembr. 1838:2036–2045.

59. Balleza, D., A. Mescola, N. Marín–Medina, G. Ragazzini, M. Pieruccini, P. Facci, and A. Alessandrini. 2019. Complex Phase Behavior of GUVs Containing Different Sphingomyelins. Biophys. J. 116:503–517.

60. Zhao, J., J. Wu, F.A. Heberle, T.T. Mills, P. Klawitter, G. Huang, G. Costanza, and G.W. Feigenson. 2007. Phase studies of model biomembranes: Complex behavior of DSPC/DOPC/Cholesterol. BBA - Biomembr. 1768:2764–2776.

61. Veatch, S.L., O. Soubias, S.L. Keller, and K. Gawrisch. 2007. Critical fluctuations in domain-forming lipid mixtures. Proc. Natl. Acad. Sci. USA. 104:17650–17655.

62. Aufderhorst-Roberts, A., U. Chandra, and S.D. Connell. 2017. Three-Phase Coexistence in Lipid Membranes. Biophys. J. 112:313–324.

63. Dymond, M.K. 2021. Lipid monolayer spontaneous curvatures: A collection of published values. Chem. Phys. Lipids. 239:105117.

64. Lentz, B.R., Y. Barenholz, and T.E. Thompson. 1976. Fluorescence depolarization studies of phase transitions and fluidity in phospholipid bilayers. 2. Two-component phosphatidylcholine liposomes. Biochem. 15:4529–4537.

65. Shah, J., J.M. Atienza, R.I. Duclos, A.V. Rawlings, Z. Dong, and G.G. Shipley. 1995. Structural and thermotropic properties of synthetic C16:0 (palmitoyl) ceramide: effect of hydration. J. Lipid Res. 36:1936–1944.

66. Chen, Z., and R.P. Rand. 1997. The influence of cholesterol on phospholipid membrane curvature and bending elasticity. Biophys. J. 73:267–276.

67. Lee, C.T., K. Venkatraman, I. Budin, and P. Rangamani. 2025. Local enrichment of cardiolipin to transient membrane undulations. Biophys. J. 124:2476–2487.

68. Giang, H., and M. Schick. 2014. How Cholesterol Could Be Drawn to the Cytoplasmic Leaf of the Plasma Membrane by Phosphatidylethanolamine. Biophys. J. 107:2337–2344.

69. Sodt, A.J., M.L. Sandar, K. Gawrisch, R.W. Pastor, and E. Lyman. 2014. The Molecular Structure of the Liquid-Ordered Phase of Lipid Bilayers. J. Am. Chem. Soc. 136:725–732.

70. Sapp, K.C., A.H. Beaven, and A.J. Sodt. 2021. Spatial extent of a single lipid’s influence on bilayer mechanics. *Phys*. Rev. E. 103:042413.

71. Lewis, R.N., and R.N. McElhaney. 1993. Calorimetric and spectroscopic studies of the polymorphic phase behavior of a homologous series of n-saturated 1,2-diacyl phosphatidylethanolamines. Biophys. J. 64:1081–1096.

72. Kučerka, N., B. van Oosten, J. Pan, F.A. Heberle, T.A. Harroun, and J. Katsaras. 2015. Molecular Structures of Fluid Phosphatidylethanolamine Bilayers Obtained from Simulation-to-Experiment Comparisons and Experimental Scattering Density Profiles. J. Phys. Chem. B. 119:1947–1956.

73. Sakuma, Y., T. Taniguchi, T. Kawakatsu, and M. Imai. 2013. Tubular Membrane Formation of Binary Giant Unilamellar Vesicles Composed of Cylinder and Inverse-Cone-Shaped Lipids. Biophys. J. 105:2074–2081.

74. Talbot, E.L., J. Kotar, L.D. Michele, and P. Cicuta. 2019. Directed tubule growth from giant unilamellar vesicles in a thermal gradient. Soft Matter. 15:1676–1683.

75. Roux, A., D. Cuvelier, P. Nassoy, J. Prost, P. Bassereau, and B. Goud. 2005. Role of curvature and phase transition in lipid sorting and fission of membrane tubules. EMBO J. 24:1537–1545.

76. Steinkühler, J., P. De Tillieux, R.L. Knorr, R. Lipowsky, and R. Dimova. 2018. Charged giant unilamellar vesicles prepared by electroformation exhibit nanotubes and transbilayer lipid asymmetry. Sci Rep. 8:11838.

77. Tsui, F.C., D.M. Ojcius, and W.L. Hubbell. 1986. The intrinsic pKa values for phosphatidylserine and phosphatidylethanolamine in phosphatidylcholine host bilayers. Biophys. J. 49:459–468.

78. Rog, T., and A. Koivuniemi. 2016. The biophysical properties of ethanolamine plasmalogens revealed by atomistic molecular dynamics simulations. BBA - Biomembr. 1858:97–103.

79. Ferraro, M., M. Masetti, M. Recanatini, A. Cavalli, and G. Bottegoni. 2015. Modeling lipid raft domains containing a mono-unsaturated phosphatidylethanolamine species. RSC Adv. 5:37102–37111.

80. West, A., V. Zoni, W.E.Jr. Teague, A.N. Leonard, S. Vanni, K. Gawrisch, S. Tristram-Nagle, J.N. Sachs, and J.B. Klauda. 2020. How Do Ethanolamine Plasmalogens Contribute to Order and Structure of Neurological Membranes? J. Phys. Chem. B. 124:828–839.

81. Girard, M., and T. Bereau. 2023. Induced asymmetries in membranes. Biophys. J. 122:2092–2098.

